# The anaerobic cryo-EM structure of the methanogenic Mtr complex reveals a nitrogenase-like [Fe_8_S_9_C] cluster bound to its active site

**DOI:** 10.64898/2026.07.03.736320

**Authors:** Tristan Reif-Trauttmansdorff, Anuj Kumar, Tomas Pascoa, Eva Herdering, Stefan Bohn, Ruth A. Schmitz, Georg K. A. Hochberg, Jan M. Schuller

**Author notes:** Corresponding author: J.M.S.

## Abstract

Methanogenic archaea conserve energy by coupling methyl-group transfer to the generation of a chemiosmotic sodium-ion (Na^+^) gradient. This central energy-conserving step is catalyzed by the membrane-bound *N*^5^-methyl-H_4_MPT:coenzyme M methyltransferase (Mtr). Here, we present high-resolution cryo-electron microscopy structures of the Mtr complex from *Methanosarcina mazei* determined under strictly anaerobic conditions. The structures reveal an unexpected, electron-dense metallocluster embedded within the central cavity of the MtrCDE trimer in the membrane plane. Based on the unique topology and density we modeled it as an [Fe_8_S_9_C] L-type cluster. It is positioned adjacent to both the coenzyme M substrate and the corrinoid cofactor of MtrA in the MtrA-MtrCDE engaged state, thereby being located right at the catalytic core of the enzyme. We could further show that binding of MtrA to MtrCDE triggers rearrangements within the interface of MtrDE that widen a putative ion-conduction pathway. The proximity of the conserved sodium-binding site to the catalytic center suggests a putative link between methyl-transfer chemistry and Na^+^ translocation. In a broader context, these findings improve our understanding of how methyl transfer, analogous to redox chemistry, can drive chemiosmotic energy conversion.

## Introduction

Methanogenic archaea are widespread in diverse environments, thrive under strictly anaerobic conditions, and catalyze the final step in the anaerobic degradation of organic matter^1^. They are the only organisms that produce methane as a primary catabolic end product and are responsible for the majority of the ∼0.5 Gt annual global net emissions of methane, a potent greenhouse gas and an important bioenergy resource^2^. Despite this global impact, the molecular mechanisms by which methanogenic archaea (methanogens) conserve energy remain incompletely understood.

In many forms of life, biological energy conservation relies on electron transport chains that couple redox reactions to ion translocation across membranes^3^. In striking contrast, methanogens employ a fundamentally different strategy, where the exergonic transfer of a methyl group between methyl-tetrahydromethanopterin (methyl-H_4_MPT), or in some species methyl-tetrahydrosarcinapterin (methyl-H_4_SPT), to coenzyme M (CoM) (ΔG°’ = −30 kJ/mol^4^), is directly coupled to vectorial sodium ion (Na^+^) translocation. These cofactors are largely restricted to methanogens^5,6^. The reaction is catalyzed by the membrane-bound N^5^-methyl-H_4_MPT:CoM methyltransferase (Mtr) and constitutes the sole energy-conserving step in hydrogenotrophic methanogenesis^7^. In methylotrophic methanogens, the reaction operates in reverse, with the Na^+^ gradient driving endergonic methyl transfer from methyl-CoM to H_4_MPT, further highlighting its central role in cellular bioenergetics^8^. The coupling of a group-transfer reaction, rather than electron transfer, to ion translocation is highly unusual. Together with ion-translocating decarboxylases in anaerobic bacteria^9^ these systems are rare examples of chemically-driven primary energy-conserving enzymes, in contrast to the more widespread redox-driven energy-conserving enzymes such as respiratory Complex I or the Rnf/Nqr complexes^10^. Despite decades of biochemical and genetic studies, the molecular mechanism by which methyl-transfer chemistry is converted into vectorial Na^+^ translocation has remained one of the major unresolved questions in bioenergetics.

The Mtr complex consists of eight subunits (MtrABCDEFGH) encoded in an operon^11^. Biochemical studies established that methyl transfer proceeds via a corrinoid-dependent two-step mechanism, in which MtrA transfers the methyl group via a methyl-Co(III)-corrinoid intermediate between H_4_MPT and CoM^4^. MtrA consists of a single transmembrane helix connected via a long flexible linker to a globular cytosolic domain, MtrA_cyt_, in which the corrinoid cofactor 5-hydroxybenzimidazolylcobamide (Factor III) is bound in a unique manner within a Rossmann fold. Structural insight of Mtr has so far been limited to a crystal structure of MtrA_cyt_^12^, a partial membrane core from *Methanothermobacter marburgensis*^13^ and a full complex from *Methanosarcina mazei* bound to the oxygen-responsive small protein MtrI^14^. However, these structures mainly defined the overall architecture of the Mtr complex, and the role of MtrI, providing only limited insights into the functioning of the complex.

Here, we determined cryo-electron microscopy (cryo-EM) structures of the Mtr complex under strictly anaerobic conditions. The structures capture conformations of MtrA_cyt_ that could approximate catalytically relevant states and reveal a nitrogenase-like [Fe_8_S_9_C]-cluster positioned at the center of the MtrCDE trimer in the membrane plane. The cluster’s location between the CoM binding site and the corrinoid cofactor suggests a role in catalysis. Conformational analysis further shows how engagement of MtrA is coupled to rearrangements in the membrane subunit MtrD that open a putative, transient ion-conduction pathway. Although the captured state contains the corrinoid in the Co(II) state and is therefore not a catalytic intermediate, it likely represents an on-pathway conformation of the catalytic cycle. Together, these findings contribute to a mechanistic understanding for methyl-transfer-driven sodium pumping, implicate a previously unrecognized role for [Fe_8_S_9_C] metalloclusters, and position Mtr as a distinct solution for biological energy conservation.

## Results

### An anoxic Mtr structure reveals a nitrogenase-like cluster in the membrane plane

To characterize the Mtr complex in a state that was not exposed to oxygen and more closely resembles a physiologically-relevant state of the enzyme, we purified it under strictly anaerobic conditions. For this, a C-terminally TwinStrep-tagged MtrE subunit was expressed in *M. mazei*, allowing rapid isolation of the complete MtrA– H complex by a single-step affinity purification from LMNG-solubilized membranes of H_2_-grown cells^14^ (Supplementary Fig. S1B,C). Under strictly anaerobic conditions, the complex displayed a dark brown color, in contrast to the light pink hue observed in all previous preparations (Supplementary Fig. S1A). We next subjected the anaerobically purified Mtr complex to redox-controlled cryo-EM, which yielded reconstructions at overall resolutions of 1.89 Å and 1.83 Å for the C1 and C3 symmetrised maps, respectively (Fig. 1A, Supplementary Fig. S2C). Notably, a high-density feature was detected at the center of the MtrCDE trimer that was absent from previously reported Mtr structures^13,14^. The high quality and resolution of the cryo-EM map enabled us to confidently model an [Fe_8_S_9_C]-cluster (Fig. 1B-E, Supplementary Fig. S5A, B), reminiscent of an L-type cluster. This is consistent with the observed metal quantification by ICP-MS, revealing 7.6 iron atoms together with 1 cobalt and 1.2 zinc ions per protomer (Supplementary Fig. S1D). L-clusters are generated in early stages of nitrogenase FeMo-cofactor biosynthesis^15^ and have been found in the FeFe-cofactor of iron-only nitrogenases^16^ (Supplementary Fig. S5C), the methylthioalkane reductase^17^, and the methyl-CoM reductase activation complex^18^. Although we do not directly observe density for the central carbide, numerous studies on nitrogenase cofactors have demonstrated that the interstitial carbide is critical for stabilization of the unique Fe–S cluster geometry and likely also contributes to the fine-tuning of catalytic properties^19–21^. Interestingly, the interatomic distances within the cluster in our model (Fe– Fe, average 2.79 Å; Fe–S, average 2.48 Å; Fe–C, 2.10 Å) are ∼5–15% longer than those reported for the FeFe cofactor^16^ (Supplementary Fig. S5B, C). This difference may reflect the distinct ligand-sphere environment of the cluster, which is highly unusual in Mtr. Rather than being stabilized via a classical side-chain coordination, the cluster appears to be accommodated within a structurally preorganized pocket (Fig. 1D, E). The binding site is formed predominantly by three cluster-supporting loops from MtrC, MtrD, and MtrE. Within these loops, proline (MtrE:Pro61, MtrE:Pro62, MtrC:Pro73) and glycine residues (MtrD:Gly47, MtrE:Gly59, MtrC:Gly71) create a compact, hydrophobic cavity that is largely devoid of bulky or polar side chains (Fig. 1D, E). This architecture is highly atypical and unique for an iron–sulfur cluster-binding site, which typically harbors coordinating residues such as cysteine or histidine to stabilize the cluster. Thus, Mtr lacks the characteristic ligation observed in canonical [Fe_8_S_9_C] clusters of nitrogenase, where the terminal Fe ions are coordinated by protein side chains or organic carboxylate ligands such as homocitrate (Supplementary Fig. S5C). Instead, the protein-proximal apical iron (Fe1, Fig. 1D, Supplementary Fig. S5B) of the cluster is associated with a tightly bound but currently unidentified ligand, which is modelled as water. Although this density could alternatively correspond to a sulfur atom, its lower peak intensity compared to the other sulfurs in the cluster, together with shorter coordination distance (2.19 Å) relative to the other Fe–S bonds, are more consistent with a water assignment (Fig. 1D; Supplementary Fig. S5B). In our structure, this apical water lies within hydrogen-bonding distance of MtrD:Gln50. In some organisms, however, this residue is replaced by methionine (Supplementary Table 1), suggesting that the weak apical interaction observed here is not universally conserved. On the opposite side of the cluster (Fe8, Fig. 1D, Supplementary Fig. S5B), facing the vestibule of the MtrCDE trimer, no well-defined apical ligand is observed. Instead, diffuse density is visible only at lower contour thresholds, suggesting structural heterogeneity (Fig. 1D). Although geometrically incompatible with direct coordination, MtrE:Asn35 is the only residue within interaction distance (2.29 Å) and may therefore provide weak polar stabilization without forming a direct bond.

**Figure 1:**
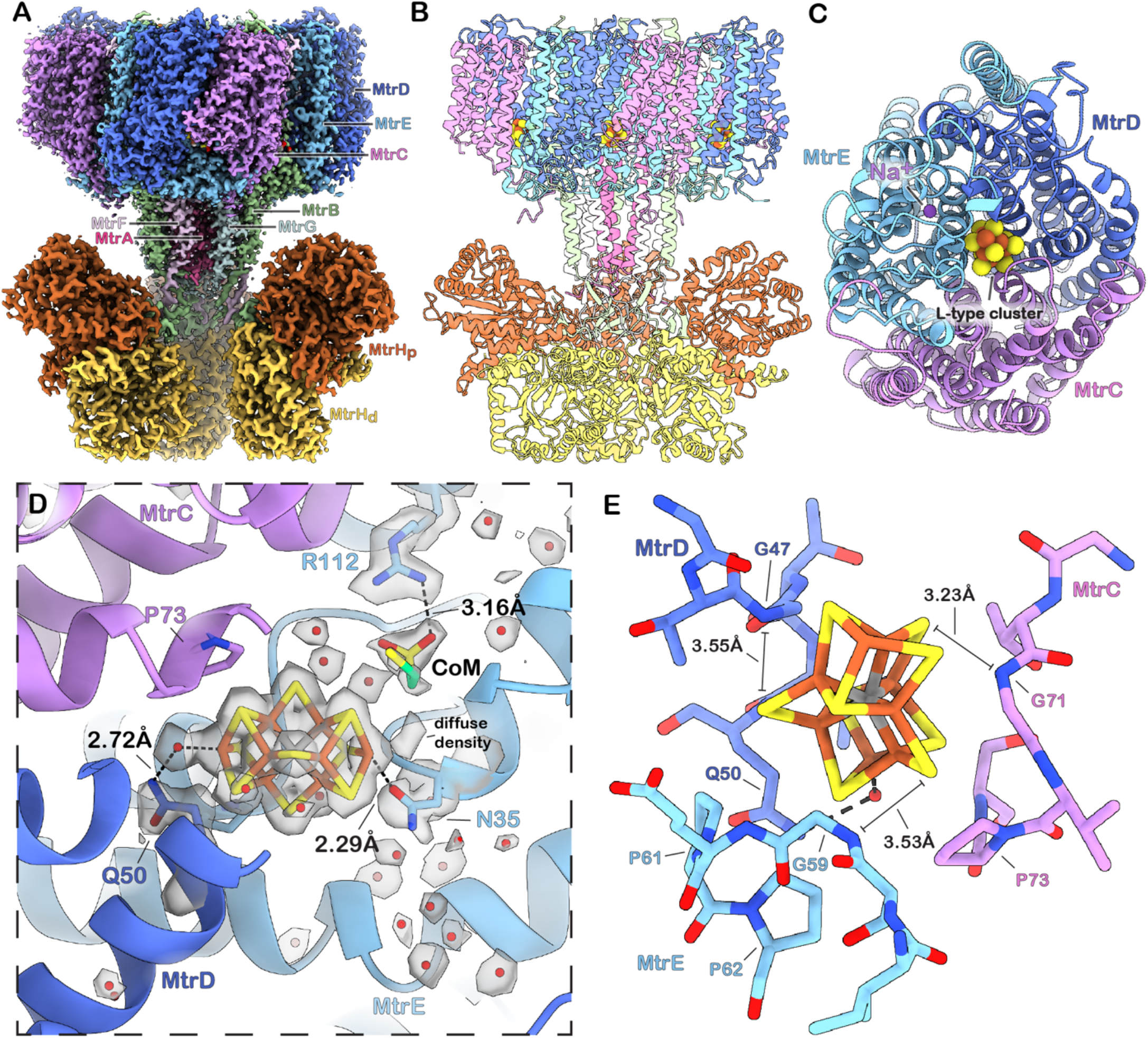
Mtr-complex binds a nitrogenase-like metallocluster at its catalytic center. **(A)** Segmented and colored cryo-EM composite (consensus) map of the Mtr core and MtrH **(B)** Corresponding atomic model in cartoon representation. **(C)** Top view of the MtrCDE trimer with the metallocluster (brown/yellow) and the Na^+^ (magenta) depicted as spheres. **(D)** Close-up view of the metallocluster with segmented consensus map around metallocluster, CoM, waters and selected residues (contour = 11.1 σ). MtrC and MtrD loops coordinating the cluster are removed for clarity. **(E)** Structural details of metallocluster binding. Three highly conserved loops from MtrC, MtrD and MtrE, each containing glycine residues, form a pocket that wedges the cluster between the belt sulfurs. Proline residues together with Gln50 form the base of the pocket on which the metallocluster rests.

Within the MtrCDE vestibule, we also observed an elongated density with a cone-shaped terminus oriented toward the guanidinium group of MtrE:Arg112. Based on its shape and agreement with the previously proposed CoM-binding site^13^, we modeled this density as CoM (Fig. 1D). Further, comparison with previously reported structures showed that the conserved Na^+^-binding site of MtrE remains unchanged. A Na^+^-ion occupies a side pocket ∼10 Å from the [Fe_8_S_9_C] cluster and is coordinated in a near-octahedral geometry by MtrE:Ser29, MtrE:Glu60, MtrE:Asp183, and three water molecules (Fig. 1C; Supplementary Fig. S4).

### MtrA_cyt_ cycles between MtrH and MtrCDE bound states

Closer examination of the cryoEM maps of the Mtr complex revealed diffuse density adjacent to the MtrCDE trimer, suggesting the presence of different conformational states of MtrA_cyt_. To resolve this heterogeneity, we performed symmetry expansion followed by focused classification, thereby identifying three distinct binding states of MtrA, two of which correspond to different stages of the catalytic cycle.

In one class, MtrA_cyt_ is bound to the membrane-proximal subunit of the MtrH methyltransferase dimer (MtrH_p_), where the methyl-group of methyl-H_4_SPT is transferred to the corrinoid-cofactor (Fig. 2A, B, E). The weaker and more diffuse density map of MtrA_cyt_ relative to MtrH_p_ indicates loose binding, consistent with its reversible association. Following local refinement, we identified the main interface contacts as MtrA:Arg103-MtrH:Glu136 and MtrA:Lys83-MtrH:Glu79, together with interactions between the corrinoid ring and MtrH:Phe156, Met200 and Pro201. Although methyl-H_4_SPT is not observed in our structure, we positioned it in the conserved binding pocket of the MtrH TIM barrel based on the binding of methyltetrahydrofolate (methyl-H4F) in the closely related methyltransferase MtgA from Desulfitobacterium hafniense^22^ to assess the positioning of the corrinoid relative to the putative substrate-binding site. This positioning was guided by fitting our MtrH structure into the electron-density map of MtgA (Supplementary Fig. S6). Notably, based on this hypothetical model, the current distance of ∼8 Å between the cobalt and the methyl-group of methyl-H_4_SPT appears too large to support direct methyl-transfer. Superimposing both structures reveals that elements of the H_4_SPT-binding site, namely loop MtrH residues 197– 205 and helix–loop elements MtrH 228–260, are displaced outward relative to the MtgA com plex. It is therefore plausible that substrate binding could induce local conformational rearrangements that allow the corrinoid cofactor to move closer to methyl-H_4_SPT, thereby positioning the cofactors in a catalytically competent geometry for methyl transfer (Supplementary Fig. S6C).

**Figure 2:**
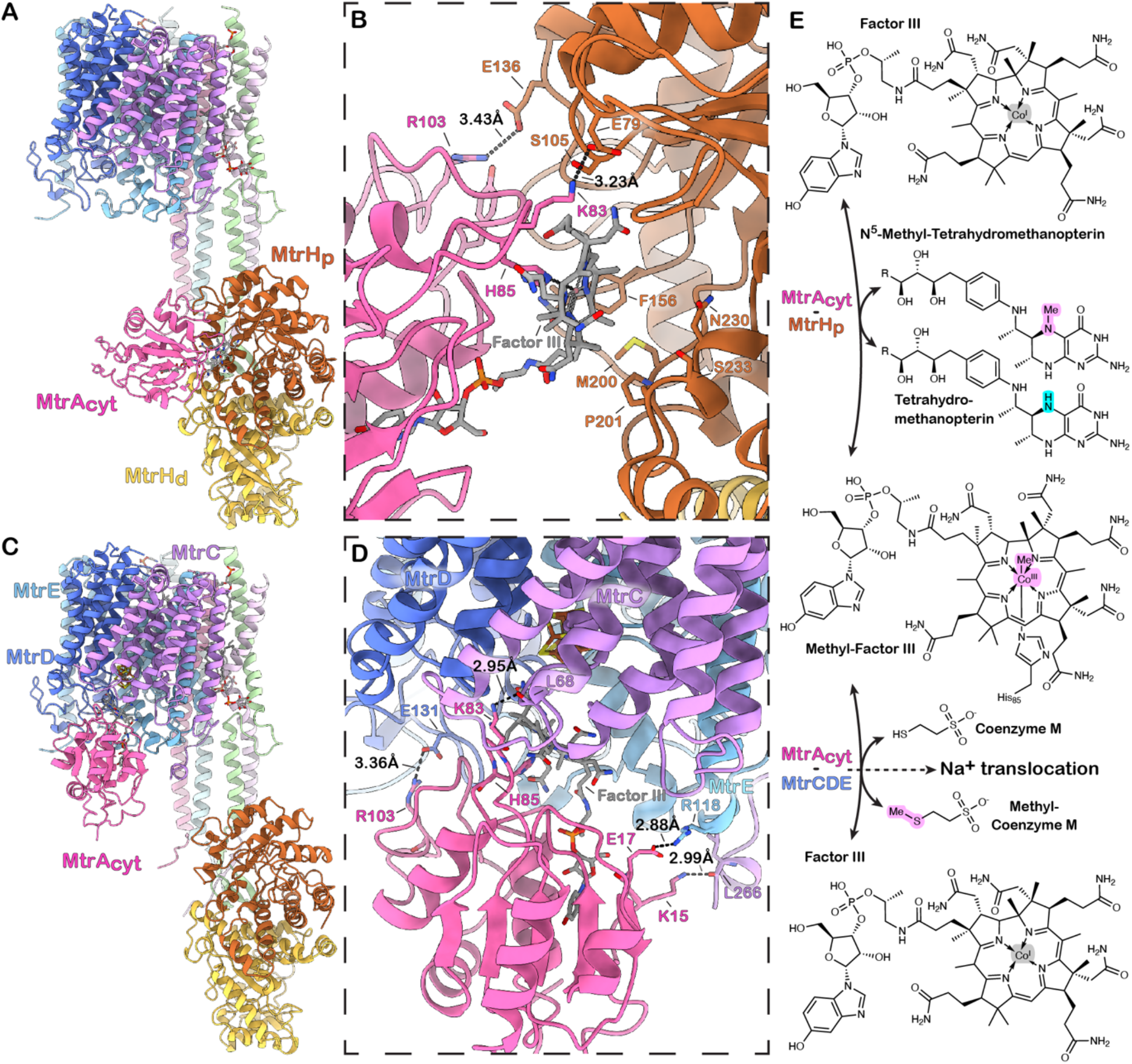
MtrA conformational states. **(A)** Side view of one protomer within the trimeric Mtr complex with MtrA_cyt_ bound to MtrH_p_. **(B)** Close-up of the MtrA_cyt_-MtrH interface in the H_4_SPT-unbound state, showing the corrinoid cofactor oriented towards the substrate-binding TIM-barrel of MtrH. **(C)** Side view of one protomer within the trimeric Mtr complex with MtrA bound to MtrCDE. **(D)** Close-up of MtrA-MtrCDE interaction. The corrinoid protrudes deeply into the vestibule harboring the CoM-binding site and the metallocluster. **(E)** Overview of the reactions catalyzed by the Mtr-complex. Atoms involved in methyl transfer, as well as the transferred methyl groups, are highlighted in the structural formulas.

We also observe a class where MtrA is positioned with its corrinoid atop Met46 of MtrE. The corrinoid-ring is further stabilized by interactions with Gln45, Met47, Phe119 (Supplementary Fig. S7). The Met-Co distance of 3.8 Å doesn’t support the formation of a covalent S-Co bond. Although a substantial fraction of the particles harbors this conformation (18% of cluster containing sites, Supplementary Fig. S2), its catalytic relevance remains unclear, as Met46 is not strictly conserved and other orthologs of Mtr can be found in which Met46 is replaced by Val, Leu or His.

Another class reveals MtrA_cyt_ positioned directly at the MtrCDE trimer. The main interface contacts include MtrA:Arg103-MtrD:Glu131, MtrA:Lys83-MtrC:L68 (carbonyl oxygen), MtrA:Glu17-MtrE:Arg118 and MtrA:Lys15-MtrC:L266 (carbonyl oxygen). In this state, the corrinoid cofactor protrudes into the vestibule containing both the metallocluster and the CoM-binding site, thereby placing the cobalt center in close proximity to the cluster and the substrate (Fig. 2C–E, Fig. 3B). This configuration is consistent with a catalytically-relevant MtrA-MtrCDE engagement state, in which the juxtaposition of the corrinoid, CoM, and the metallocluster raises the possibility that the cluster participates directly in catalysis, for example by contributing to CoM activation.

**Figure 3:**
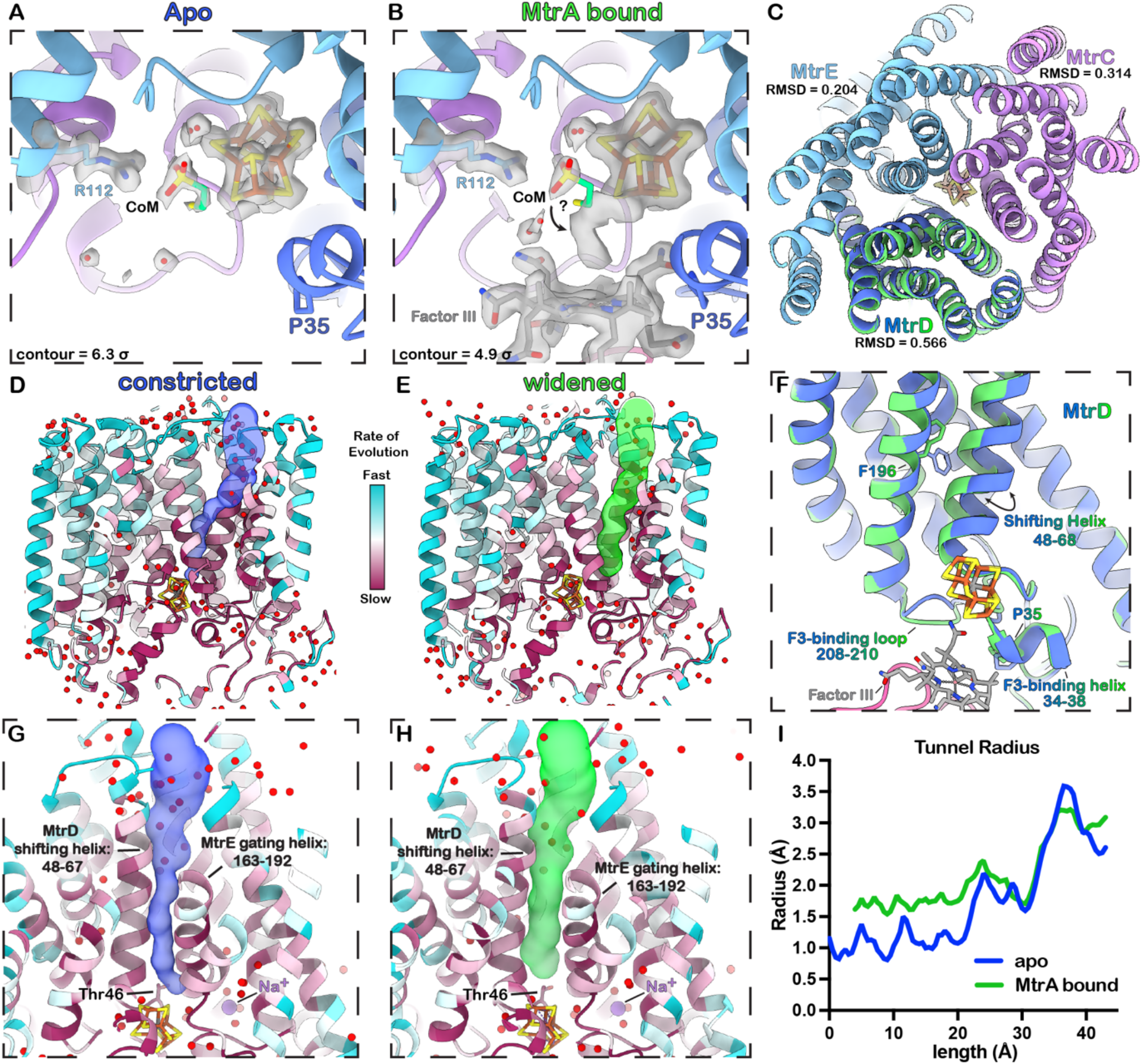
MtrA binding triggers widening of a putative ion-conduction pathway. **(A, B)** Close-up of the corrinoid-binding site at the MtrCDE vestibule in the apo and MtrA-bound states. Corrinoid binding correlates with ambiguous density extending from the apical iron of the metallocluster, suggesting alternative substrate states. **(C)** Superposition of the apo and MtrA-bound MtrCDE trimers with pairwise Cα RMSD values indicated. To highlight the helix shift, MtrD is shown in dark blue in the apo state and green in the MtrA-bound state. **(D, E)** Side views of MtrCDE in the constricted (apo) and widened (MtrA-bound) states colored by evolutionary rate. Predicted tunnels at the MtrD–MtrE interface are shown as transparent surface representations. **(F)** Close-up of the superposition of MtrD from the apo (dark blue) and MtrA-bound (green) states, highlighting structural rearrangements. **(G, H)** Tilted top views of the MtrD–MtrE interface in the constricted and widened conformations, showing predicted tunnels and Thr46 used as the prediction starting point. **(I)** Tunnel radius profiles of the apo and MtrA-bound states plotted against tunnel length. As the apo tunnel is longer and originates further downstream, profiles were aligned using overlapping regions in the second half.

### MtrA binding to MtrCDE positions the corrinoid in proximity to the cluster

Because MtrA_cyt_ binding to MtrCDE remained heterogeneous within this state, we carried out additional 3D variability analysis of the MtrA_cyt_–MtrCDE subcomplex. The analysis revealed a dominant mode of conformational heterogeneity consistent with progressive insertion of MtrA_cyt_ into the MtrCDE vestibule, accompanied by increased definition of the corrinoid density in the tightly engaged state (Movie 1, 2). Refinement of the tightly bound conformation resolved a ligand-associated density extending between the apical iron of the metallocluster and the cobalt center of the corrinoid cofactor. The cobalt appears to be in the Co(II) state, as indicated by the absence of an upper axial ligand and the presence of the lower axial ligand His85 of MtrA. (Supplementary Fig. S4, Factor III, MtrA-MtrCDE core map).

This composite density cannot be modelled unambiguously and instead points to a mixture of CoM binding states (Supplementary Fig. S8). Although corrinoid insertion into the vestibule correlates with changes in substrate binding, it remains unclear how the observed, non-catalytic Co(II) state contributes to this arrangement or how it differs from a methylated corrinoid or tetracoordinate Co(I) state.

### MtrA binding remodels the membrane domain and opens a potential sodium translocation pathway

Further examination of the 3D variability analysis of the MtrA_cyt_–MtrCDE subcomplex revealed that the insertion of the corrinoid within the vestibule is accompanied by a concerted widening and local rearrangements of the transmembrane MtrCDE trimer. (Movie 3). Comparison of the apo and MtrA-bound MtrCDE subcomplexes shows that MtrC and MtrE remain rather unchanged but MtrD displays more substantial rearrangements (Figure 3C). These consist of small movements of the F3-binding loop (residues 208–210) and Factor-III-binding helix (residues 34–38), together with a pronounced outward displacement of transmembrane helix TM2 (residues 48–68) (Fig. 3F). Notably, TM2 is the only transmembrane element showing pronounced conformational variability, suggesting it could play a potential role in gating or ion translocation.

To evaluate whether the rearrangements cause changes in a putative pathway through the membrane domain, we performed computational tunnel analysis using CAVER^23^. Because MtrD:Thr46 is positioned directly above the metallocluster and at the base of the shifting helix, we used this residue as the starting point. In the MtrA-bound state, the analysis identified a tunnel originating near the base of the shifting helix, traversing the membrane region along the MtrDE interface between TM2 of MtrD and TM5 of MtrE, and opens into a water filled funnel at the extracellular site (Fig. 3E, H; Supplementary Fig. S10B, C and S11). In contrast, in the apo state, the predicted tunnel is highly constricted, with TM2 of MtrD and TM5 of MtrE forming a hydrophobic constriction along the MtrDE interface (Fig. 3D, G).

Comparison of the tunnel profiles shows that the apostate tunnel contains multiple constrictions with radii below 1 Å, reaching a minimal radius of 0.81 Å. By contrast, the MtrA-bound state features a continuous tunnel with a minimal radius of 1.54 Å (Fig. 3E).

Thus, although the observed state is unlikely to support Na^+^ passage, the MtrA-induced rearrangements strongly implicate this pathway in ion translocation (Movie 4, 5).

### Conservation analysis supports proposed pathway at the MtrDE interface

To get independent support for the functional relevance of structural features of the MtrCDE trimer, particularly the pathway at the MtrDE interface, we performed evolutionary rate analysis. We reasoned that sites involved in the sodium pathway would be expected to evolve more slowly than other sites. To measure this, we inferred a concatenated maximum likelihood phylogeny of MtrCDE using a substitution model that allows for site specific evolutionary rates. We then extracted these rates and projected them onto the structure by color mapping (Fig. 3D, E, G, H; Supplementary Fig. S9-11). This analysis revealed that rapidly evolving regions are predominantly at the membrane-exposed surface of MtrD and MtrC, as well as at the extracellular regions of the trimer (Supplementary Fig. S9A, B). In contrast, the MtrA binding site, the cytosolic vestibule and partial regions of MtrE that form the interface with the stalk exhibit slower evolutionary rates indicating strong conservation (Supplementary Fig. S9C, D). Analysis of the individual MtrCDE subunits further shows that both the center of trimerization and the metallocluster binding sites are highly conserved. Interestingly, we see a strong conservation of residues lining the channel at the MtrDE interface, comparable to that observed at the metallocluster binding site. In contrast, the corresponding, pseudo-symmetric interface forming regions of MtrCE and MtrCD show substantially lower conservation (Supplementary Fig. S10, S11). Together with the structural comparison and tunnel analysis, these observations support the functional relevance of a conserved pathway at the MtrDE interface, adjacent to the sodium binding site, that appears to be dynamically regulated by conformational changes associated with MtrA binding.

## Discussion

In this work, we identify an unexpected nitrogenase-like [Fe_8_S_9_C] metallocluster bound within the membrane plane of the methanogenic Mtr complex. Importantly, the cluster is located within the MtrCDE trimer directly adjacent to the CoM binding site and close to the corrinoid cofactor of MtrA as well as a highly conserved sodium binding site. Although the present structures do not define the underlying chemistry in atomic detail, they place the corrinoid, substrate, and metallocluster in a configuration compatible with direct participation of the cluster in catalysis.

In a broader context, our structures provide a framework that unifies decades of biochemical work on the Mtr complex and helps explain several previously reported observations. Earlier work established that Mtr catalyzes the exergonic transfer of a methyl group from *N*^5^-methyl-H_4_MPT to CoM through a corrinoid-dependent two-step reaction, that catalysis requires the super-reduced Co(I) state of MtrA, and that methyltransfer to CoM is strongly stimulated by Na^+24,25^. Reconstitution experiments further demonstrated that Mtr functions as a primary Na^+^ pump, translocating between 1-2 Na^+^ per methyl group transferred, while mechanistic models proposed that Na^+^ pumping is triggered during the sodium-dependent demethylation of the MtrA corrinoid^26^. Furthermore, an earlier study also reported that Mtr contains eight non-heme iron atoms and eight acid-labile sulfur atoms per protomer^27^. EPR experiments with Mtr yielded spectra consistent with a 4Fe–4S cluster, and treatment with ferricyanide led to complete inactivation^28^. However, these findings were generally assumed as arising from contamination by co-purifying polyferredoxin complexes, particularly because no conventional FeS-cluster-binding site could be identified in the sequence of Mtr^29^. Instead, a catalytic Zn^2+^ was proposed to serve as a Lewis acid to activate the HS-CoM for a nucleophilic attack on the methyl-corrinoid^29^. Our structures now reconcile these observations by revealing that Mtr contains an L-type [Fe_8_S_9_C] cluster, whose architecture would not have been predicted from conventional FeS-binding motifs.

The data support a model in which the metallocluster contributes to the second half-reaction of catalysis, during which the methyl group is transferred from methyl-Co(III)-corrinoid to HS–CoM. In the catalytically engaged state, the corrinoid inserts deeply into the MtrCDE vestibule and is positioned immediately adjacent to both the CoM binding site and the apical iron of the [Fe_8_S_9_C] cluster. Additional density extends between the corrinoid cobalt and the apical iron. Although this density cannot be unambiguously assigned, its position and shape is compatible with CoM- and/or methyl-CoM-containing states located directly at the cluster. Importantly, the observed corrinoid appears to adopt a non-catalytic Co(II) state. Although already providing a valuable structural snapshot, the organization of the active site in the catalytically relevant methylated Co(III) or axial-ligand-free Co(I) states awaits further characterization.

At minimum, these observations indicate that the cluster forms part of the chemical environment in which methyl transfer occurs. A plausible interpretation is that the apical iron contributes to substrate binding and thiol activation, thereby promoting nucleophilic attack by HS–CoM. Such catalytic involvement of the metallocluster is not without precedent, as iron–sulfur cofactors can participate directly in substrate binding and activation rather than solely in electron transfer. Aconitase provides a classical example, as its substrate citrate binds directly to the unique iron of the [4Fe–4S] cluster during catalysis^30^. By contrast, thiol activation in corrinoid-dependent methyltransferases is more commonly mediated by Zn^2+^, as in cobalamine dependent methionine synthase (MetH) and MtaA from the methanol:CoM-methyltransferase system^31,32^. Although ICP-MS analysis revealed approximately one zinc atom per Mtr protomer, a corresponding zinc-binding site is not apparent, particularly near both CoM and the corrinoid. Why Mtr would employ a large iron–sulfur metallocluster instead of a zinc center is not yet clear. One possible explanation lies in the flexible coordination chemistry of iron, particularly within iron–sulfur clusters as in aconitase, where it can accommodate coordination numbers greater than four.

This adaptable coordination environment may increase ligand flexibility, while the comparatively loose binding mode of the cluster may allow it to shift and rotate within its binding site. In the context of coupling local substrate chemistry to large-scale conformational changes, these characteristics might provide a critical functional advantage.

However, how local chemistry at the L-cluster is coupled to Na^+^ release is not resolved by our structures. Nevertheless, comparison of the different structural states suggests a plausible working hypothesis. The most direct connection between the metallocluster and the Na^+^ binding site is mediated by MtrE:Asn35. In the apo structure, Asn35 is positioned adjacent to the diffuse density connected to the vestibule-facing apical iron and is linked, via a bridging water molecule, to Asp183, a key ligand of the Na^+^-binding site. How Asn35 might relay events at the iron center to the sodium site remains unclear. One possibility is that close to the methyl-transfer transition state, the Na^+^ binding site is destabilized, promoting movement of Na^+^ toward and past the cluster, transiently coordinated by conserved MtrD:Thr46, MtrE:Ser179, and backbone carbonyls, before entering the extracellular tunnel (Fig. 4).

**Figure 4.**
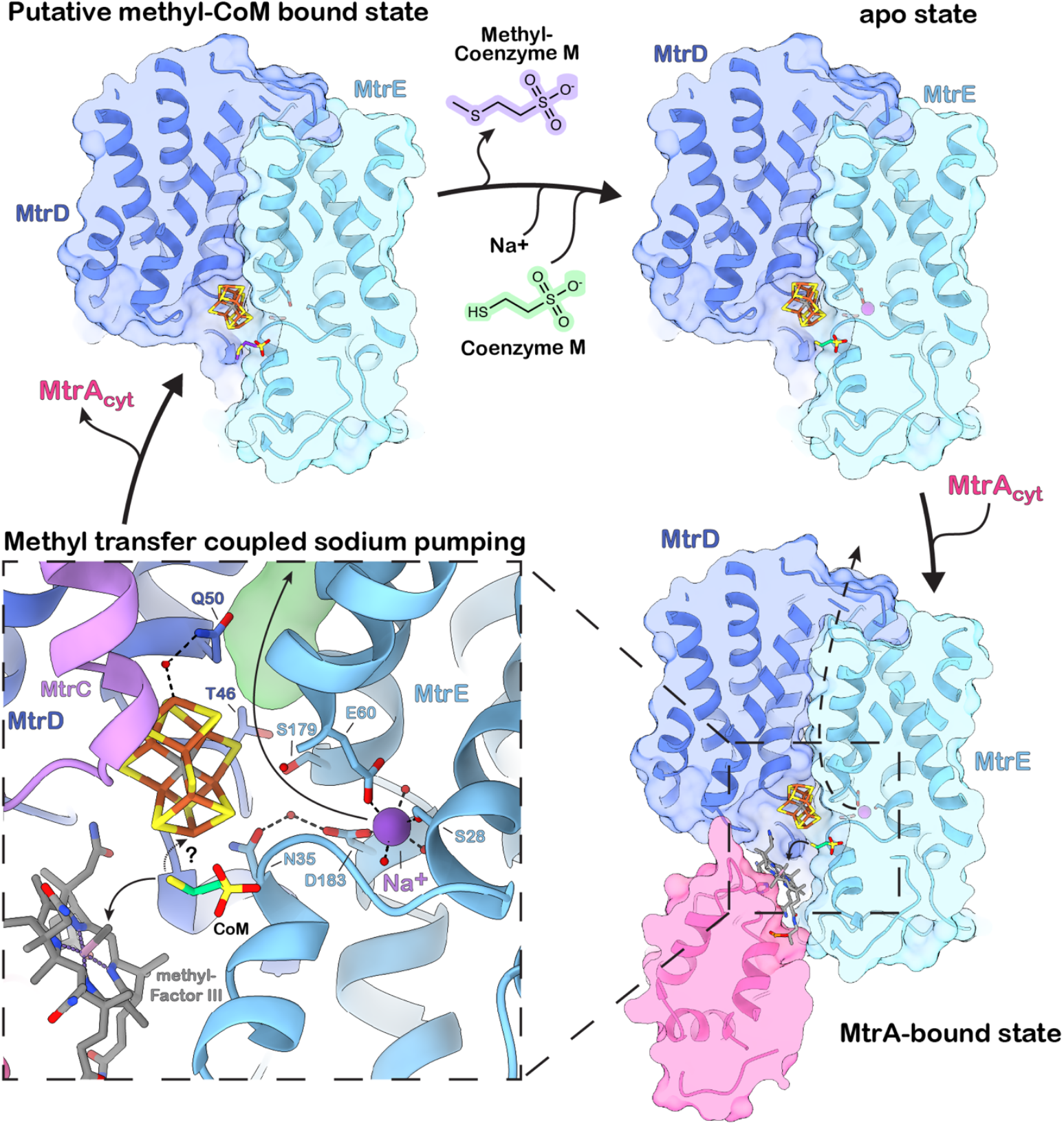
The L-cluster is positioned between the methyl transfer and the sodium translocation sites. The apo state is consistent with an inward-open state that transitions into an inward-occluded state accompanied by priming of the outward-open state upon MtrA-binding. The potentially high-energy outward-open conformation may be reached transiently near the transition state of the methyl transfer reaction.

Overall, the structural data further support a previously proposed alternating-access mechanism^13,33^, a transport principle ubiquitous among membrane transporters. One gating element of Mtr appears to be the MtrDE interface, which comprises a relatively rigid component (e.g. MtrE:TM5) and a mobile component (e.g. MtrD:TM2) that moves relative to it. In this framework, the apo Mtr structure is compatible with an inward-open state in which the sodium-binding site accessible from the cytosol. Binding of MtrA–MtrCDE and insertion of the corrinoid into the vestibule block cytosolic access and trigger conformational changes that widen a putative sodium-translocation pathway between MtrD and MtrE. In the present structures, the sodium ion is still bound strictly within its binding site, and no fully outward-open state is observed. MtrA binding is therefore unlikely to be sufficient to trigger sodium translocation, and this conformation likely represents an inward-occluded state that precedes the outward-open state, which may only be transiently populated during methyl transfer.

More broadly, these findings place Mtr alongside Complex I and the Rnf/Nqr complexes within a common conceptual framework of ion-translocating bioenergetic machines, however with a unique driver of Na^+^ transfer. In Complex I, local redox chemistry is coupled to proton translocation through long-range conformational changes^37^, whereas in Rnf and Nqr, electron transfer through flavin and iron–sulfur cofactors is linked to Na^+^ transport by membrane-embedded redox modules that regulate ion-binding sites and access pathways^38^. Recent structural work on Rnf further identified a membrane-embedded [2Fe–2S] cluster that may participate directly in coupling by attracting Na^+^ and triggering conformational changes that alter membrane accessibility^39,40^. Against this background, Mtr represents a distinct manifestation of the same overarching principle. In this system, however, it is not electron transfer but methyl-group transfer that drives vectorial sodium pumping.

## Materials and Methods

### Chemicals

Unless specified otherwise, chemicals were acquired from Sigma-Aldrich, Carl Roth GmbH + Co. KG, Roche and Serva GmbH.

### Strains and plasmids

MtrE was expressed with a C-terminal TwinStrep tag (TS-tag) in *Methanosarcina mazei* DSM 3647 using plasmid pRS1743 as described previously^14^. Briefly, pRS1743 consists of the backbone pRS1595 (a shuttle vector for *M. mazei* and *Escherichia coli)*^41^, harbouring the puromycin resistance gene and the *E*.*coli* and methanogenic replicons. It further contains the constitutive *pmcr*B promotor, including a ribosome-binding site driving expression of C-terminally TS-tagged *mtr*E gene (*MM_1547*).

### Cultivation and harvest

*M. mazei* Gö1 was grown at 37 °C under an H_2_/CO_2_ gas mixture (80%/20% ) as the growth substrate, in an orbital shaking incubator at 100 rpm in closed, anoxic glass tubes in volumes ranging from small volumes (e.g., 5 mL in Hungate tubes) to larger volumes of up to 1 L in 2 L Duran bottles. The medium used was similar to DSMZ120 but buffered with 20 mM PIPES (pH 7.0) instead of NaHCO_3_ and puromycin was added at a final concentration of 2-5 mg/L (Alomone Labs). Media were prepared aerobically and turned anoxic after autoclaving by repeatedly cycling the gas-phase with N_2_. Complete anoxic conditions were achieved by addition of 40 mg/L of Na_2_S followed by visually monitoring the reduction of resazurin. Vitamins, minerals and antibiotics were added after autoclaving. Cells were harvested during late log-phase at an OD_600_ of 0.7-1 at 10 000 x g at 4 °C. To achieve an anoxic harvest, centrifugation bottles and lids were pre-incubated for at least 16 h under strict anoxic conditions inside of a vinyl anaerobic chamber with 95-98% N_2_ and 2-5% H_2_ (Coy Laboratory Products). Cultures were shuttled into the chamber and successively transferred into the centrifugation bottles. Tightly sealed bottles were moved outside the tent for centrifugation. Resazurin in the growth media served as an indicator to confirm maintenance of anoxic conditions throughout the process. Harvested cells were stored at -80 °C.

### Purification of the Mtr complex via TwinStrep-tagged MtrE

Protein purification was conducted as described previously^14^, with all steps being conducted under strict anoxic conditions inside of an anaerobic chamber with 95-98% N_2_ and 2-5% H_2_. Briefly, cells were suspended in Buffer A (50 mM MOPS/NaOH pH 7.0, 10 mM MgCl_2_, 150 mM NaCl, 2mM DTT) and lysed by sonication. After clarification by centrifugation (20,000 x g) membrane solubilization was performed directly in the lysate with 1.5% LMNG for 12–16 h at 4 °C. The solubilized lysate was incubated with Strep-Tactin Sepharose resin, washed with Buffer A, and eluted with Buffer A containing 2.5 mM desthiobiotin. The protein was subsequently concentrated using 100 kDa centrifugal filters.

### Inductively coupled plasma mass spectrometry (ICP-MS) analysis

To quantify the metal content in purified protein fractions, concentrated nitric acid (67% w/v HNO_3_; Chemsolute, superpure grade) was added to the protein solution to achieve a final concentration of 11% HNO_3_. Samples were then mineralized at 80 °C for 4 hours and subsequently diluted with Chelex-treated water to a final concentration of 2% HNO_3_, yielding a total volume of 250 µL per sample. Internal standards, indium (2 ppb) and germanium (20 ppb) (both from VWR Merck), were added to each sample. Elemental analysis was conducted using an inductively coupled plasma mass spectrometer iCAP Q (ICP-MS, Thermo Fisher Scientific), equipped with an SC4DX autosampler (Elemental Scientific) and a MicroFlow PFA-100 nebulizer. An external calibration was performed using the ICP multi-element standard solution XVI (VWR Merck). Samples were introduced via a peristaltic pump and analyzed for Ca^44^, Fe^56^, Co^59^, Ni^60^, Zn^66^, Mo^95^ and W^182^. The ICP-MS operated with a reaction cell using a helium/hydrogen gas mixture (93/7%) at 5 mL min^−1^, an argon carrier gas at 0.72 L min^−1^, and an argon plasma makeup gas flow of 1 L min^−1^. Data acquisition was performed in triplicate using Qtegra software v2.18 (Thermo Fisher Scientific). Blank values and quality thresholds were calculated using protein-buffer standards. The measured concentrations (initially in ppb) were converted to molar units (µM) of metal per sample for quantitative analysis.

### Grid preparation

For redox-controlled cryo-EM of the Mtr complex grids (QUANTIFOIL R 2/1, copper 300 mesh grids (Quantifoil Micro Tools)) were prepared using a Vitrobot Mark IV (Thermo Fisher Scientific) placed in a vinyl anaerobic chamber with 95-98% N2 and 2-5% H2 (Coy Laboratory Products). Prior to sample application, the grids were glow-discharged for 25 seconds at 15 mA using a PELCO easiGlow device (Ted Pella). Purified Mtr at a concentration of 10 mg/mL was pre-mixed in a 9:1 ratio with 10 mM CHAPSO, resulting in a final CHAPSO concentration of 1 mM, to counter preferred particle orientation. Immediately after mixing, 4 μL of the protein solution was applied to the grid and plunge-frozen in liquid ethane. Grids were blotted for 8 seconds with a blot force of 8, while the chamber-atmosphere being kept at 4 °C and 100% humidity.

### Cryo-EM data collection and processing

Cryo-EM data for anaerobically preserved Mtr was acquired using Smart EPU Software on a TFS Krios G4 cryo-TEM operating at an accelerating voltage of 300 keV and equipped with a Falcon 4i Direct Electron Detector (Thermo Fisher Scientific) and Selectrics Energy Filter set at a 10 eV slit width. Data were acquired at a nominal magnification of 165 000 x corresponding to a calibrated pixel size of 0.73 Å. Images were acquired with an exposure dose of 55 e^−^ Å^-2^ in counting mode and exported as an electron-event representation (EER) file format. Three different datasets were acquired containing 4957 (Batch 1), 15730 (Batch 2) and 13672 (Batch 3) micrographs respectively (Supp. Fig. 1A). The entire processing was done in CryoSPARC^1^ v.4 and v.5 BETA (Supp. Fig. 1C). The three batches were initially processed separately, and curated particles from each batch were later pooled for joint processing. The EER files were fractionated into 60 frames, motion corrected using patch motion correction^43^, followed by contrast transfer function (CTF) estimation. For Batch 1, acquired more towards the center of the grid-holes and showing a more uniform particle distribution, a Topaz model was trained on 500 manually picked particles, followed by Topaz extract, yielding 154,286 particles. For Batch 2 and 3, acquired near the grid-hole edges and showing tighter particle packing, blob picker was used yielding 3,002,668 and 1,786,390 particles, respectively. Particles were extracted at a box size of 576 pixels. All Batches were subjected to heterogenous refinement to sort particles in 3D using an initial map, that was generated from a testing dataset as an input seed. Batch 1 was heterogeneously refined using five input maps. Three refinements were subsequently pooled (136,924 particles), grouped into 35 exposure groups after beam-shift import, subjected to reference-based motion correction (RBMC), and refined using non-uniform refinement, yielding a resolution of 1.92 Å, or 1.83 Å with C3 symmetry applied.

Batch 2 was heterogeneously refined in three sub-batches using six input maps. The best refinements of each sub-batch were pooled (850,619 particles), grouped into 37 exposure groups after beam-shift import, subjected to RBMC, and refined using non-uniform refinement, yielding a resolution of 1.96 Å, or 1.91 Å with C3 symmetry applied. This was followed by a second round of heterogeneous refinement using 6 input maps for further clean-up. Three of these refinements (603,731 particles) were further non-uniform refined to 1.96Å. Batch 3 was heterogeneously refined in one batch using six input maps. The best two refinements (682,517 particles) were subjected to a second round of heterogenous refinement using 6 input maps. The best refinement (389,110 particles) was grouped into 37 exposure groups after beam-shift import, subjected to reference-based motion correction (RBMC), and refined using non-uniform refinement, yielding a resolution of 1.95 Å. Merging the final particles from Batch 1, 2 and 3 (1,125,359 particles) was followed by non-uniform refinement yielding 1.89Å or 1.83 Å with C3 symmetry applied.

#### Symmetry expansion

We performed symmetry expansion, effectively tripling the number of asymmetric units (here called sites), enabling more exhaustive analysis of conformational differences within a protomer. To focus on sites containing the iron-sulfur-cluster we started off with focused 3D classification without alignment into two classes using a focus mask covering the cluster binding pocket and a resolution filter of 5 Å, yielding 1,674,550 sites with metal-cluster and 1,701,517 sites without cluster. Successive processing was performed solely with the metal-cluster-containing sites.

#### Consensus map

Processing of the consensus map was initiated with 3D classification into 15 classes using a mask covering the cluster-containing MtrCDE sub-complex and a 3 Å resolution filter to further remove sites containing a destroyed or partially destroyed metal-cluster. The best classes (1,004,563 sites) were homogenously reconstructed to yield a resolution of 1.89Å. To further resolve the flexible, dimeric MtrH on each trimeric site, three local refinements were performed using masks, each covering a different MtrH dimer, yielding three MtrH dimer maps at resolutions of 2.14 Å, 2.29 Å and 2.29 Å respectively. In ChimeraX different commands were used to generate a composite consensus map. *vop scale* was used to adjust the map thresholds, *fit in map* and *vop resample* were used to properly align the refinements and *vop max* was applied to the homogenous reconstruction of the Mtr core and the three MtrH dimers to generate a composite map.

#### MtrA_cyt_ conformational states

To initiate the analysis of the different dynamic states of MtrA_cyt_, focused 3D classification into six classes at 5 Å resolution using a mask covering the dynamic range of MtrA_cyt_ movement was performed with the metal-cluster-containing sites. The resulting classes were further used to resolve MtrA_cyt_ bound to MtrCDE, MtrA_cyt_ bound to MtrH and MtrA_cyt_ bound to MtrE:M46.

#### MtrA_cyt_ at MtrCDE

The corresponding class (309,078 sites) showing MtrA_cyt_ at MtrCDE was 3D classified into two classes using a mask covering the cluster-containing MtrCDE sub-complex and a 3 Å resolution filter to further remove sites containing a destroyed or partially destroyed metal-cluster. The best class was subjected to 3D variability analysis at a filtered resolution of 3 Å with three modes using a loose mask covering MtrA_cyt_CDE. The resulting three components were displayed in simple and further in cluster mode using 8 clusters. While two components were dominated by noise the third displayed strong MtrA_cyt_ variability with MtrA_cyt_ showing conformations ranging from tightly to loosely bound at the MtrCDE trimer reflecting a continous insertion movement of MtrA_cyt_ at its binding site (Movie 1). Pooling and homogenous reconstruction of the two clusters (28,603 sites) showing MtrA_cyt_ most tightly bound yielded a map of 2.16 Å resolution. Together with a local refinement of MtrH (3.49 Å) a composite map was constructed in ChimeraX as described above.

#### MtrA_cyt_ at MtrH

Four 3D classes showing MtrA_cyt_ bound to neither MtrCDE nor MtrE:M46 were pooled (1,067,739 sites) and 3D classified into ten classes using a mask covering the MtrH dimer and MtrA_cyt_ bound to MtrH (MtrAH mask). Selected classes were pooled (848,345 sites) and locally refined using the MtrAH mask before being subjected to another round of 3D classification into six classes at 6 Å using the MtrAH mask. The class showing MtrA bound to MtrH (142,972 sites) was further subjected to 3D Variability analysis (5 Å filter, three modes) using the MtrAH mask. Clusters with well-resolved MtrAcyt (93,115 sites) were pooled and locally refined to 2.51 Å using the MtrAH mask. Using a mask covering MtrA_cyt_ bound to MtrH we locally refined MtrA_cyt_ to a resolution of 3.38Å. Further, using these sites, we generated a homogenous reconstruction yielding the Mtr core at 2.06 Å. Subsequentially, we generated a composite map in ChimeraX as described above using the local refinements of MtrAH and MtrA_cyt_ and the homogenous reconstruction of the core.

#### MtrA_cyt_ at MtrE:M46

The corresponding class showing MtrA_cyt_ bound to MtrE:M46 was 3D classified into three classes at a resolution of 5 Å using a mask covering MtrA_cyt_. The best class (98,911 sites) was homogenously reconstructed to a resolution of 2.07 Å. To resolve MtrA_cyt_ we performed particle subtraction using a Mtr-consensus mask covering the Mtr core together with the three MtrH dimers. The subtracted particles were subjected to local refinement using a mask covering MtrA_cyt_ yielding at map at 3.64 Å. Subsequentially, using the homogenous reconstruction of the core and the local refinement of MtrA_cyt_ we generated a composite map in ChimeraX as described above.

### Model building and refinement

PDB:9QTS was used as an initial model. ChimeraX and Coot (v.0.9.8.91)^44^ were used for manual model building. The presence of water molecules was predicted using douse, as implemented in the PHE-NIX package (v.1.21-5207), with standard settings followed by a manual curation in coot. Real-space refinements of models were performed iteratively with the PHENIX. Ramachandran, reference model and secondary structure restraints were applied during refinement. CIF files for ligands like the [Fe_8_S_9_C]-Cluster were created using AceDRG in the CCP4 suite or eLBOW in PHENIX. Maps were graphically depicted using ChimeraX and Coot.

### Tunnel prediction

Tunnel calculations were performed using the CAVER 3.0.3 PyMOL plugin. The apo and MtrA-bound structures were used as input models, with the Cβ atom of MtrD:Thr46 defined as the starting point. Calculations were performed using a minimum probe radius of 0.6 Å for the apo structure and 0.8 Å for the MtrA-bound structure, a shell depth of 10 Å, a shell radius of 10 Å, and 12 approximating spheres. The desired starting radius and maximum starting distance were set to 2 Å and 4 Å for the apo structure, and 5 Å and 5 Å for the MtrA-bound structure, respectively. Waters, lipids, CoM, and sodium ions were omitted from the calculations.

### Evolutionary rate analysis

We collected a dataset of sequences of MtrCDE spanning archaeal species diversity, using the sequence similarity search function of the Alphafold database. Our dataset contains only species for which we identified copies for all three genes. We then aligned these sequences using Mafft version v7.520^45^ using default settings and trimmed the sequences manually such that the sequences from *M. mazei* were gap free (i.e. we trimmed out all insertions that were not present in the *M. mazei* proteins). We then inferred a maximum likelihood phylogeny using IQtree2^46^. The best fit model was determined to be LG+F+R7 using IQtrees model finder procedure. We extracted the site-specific rates using the -rate flag and used the Posterior mean site rate weighted by posterior probability for downstream analysis. The advantage of this method relative to conventional conservation analyses is that it is not sensitive to biased sequence sampling, because it takes into account the tree structure when rates are inferred. The rate model used here also takes into account what kinds of substitutions (conservative vs non-conservative) occur at each site across the phylogeny.

## Data availability

The cryo-EM maps generated in this study have been deposited in the Electron Microscopy Data Bank (EMDB). Mtr apo structure: Composite map - EMD-58097, Consensus map – EMD-58079, local ref. MtrH.1 – EMD-58094, local ref. MtrH.2 – EMD-58081, local ref. MtrH.3 – EMD-58084. MtrA bound to MtrCDE structure: Composite map EMD-58098, Consensus map – EMD-58092, local ref. MtrH – EMD-58095. MtrA bound to MtrH structure: Composite map – EMD-58111, Consensus map – EMD58078, local ref. MtrH – EMD-58107, MtrA bound to MtrH EMD-58080. MtrA bound to MtrE:M46 structure: Composite map - EMD-58076, Consensus map – EMD-58074, MtrA bound to MtrE:M46 – EMD58075. Corresponding coordinate files for the composite maps have been deposited in the RCSB Protein Data Bank (PDB). Mtr apo structure: 30VS. MtrA bound to MtrCDE structure: 30VT. MtrA bound to MtrH structure: 30WF, MtrA bound to MtrE:M46 structure: 30UY. Coordinate files for MtgA are accessible under the accession code 6SJN. Coordinate files for Mtr (*M. marburgensis*) are accessible under the accession code 8Q3V.

## Acknowledgments

We acknowledge the cryo-EM Platform (CEMP), Helmholtz Munich, for data acquisition. We thank Rolf Thauer for continuous support and helpful discussions. We thank Duncan Kountz for critical reading of the manuscript and for valuable comments and suggestions that improved the study. We thank Darja Deobald from the Helmholtz Centre for Environmental Research (UFZ), Leipzig, Germany, for performing metal quantification through ICP–MS. This work was funded by the Deutsche Forschungsgemeinschaft (DFG, German Research Foundation) under Germany’s Excellence Strategy – EXC 3048 – Project number 533620160: Microbes-for-Climate (M4C) Cluster of Excellence, Synmikro, Marburg, Germany and the European Union’s Horizon 2020 research and innovation program under grant agreement No 101075992 (Two-CO2-One). The views and opinions expressed are those of the author(s) only and do not necessarily reflect those of the European Union or the European Research Council. Neither the European Union nor the granting authority can be held responsible for them. This paper was typeset with the bioRxiv word template by @Chrelli: www.github.com/chrelli/bioRxiv-word-template

## Author contributions

T.R.T. cultivated cells, isolated and characterized the complex and vitrified cryo-EM grids. A.K. and S.B. performed the initial grid screening and cryo-EM data collection. T.R.T. conducted the cryo-EM data analysis, model building, and refinement. T.R.T., T.P., A.K., and J.M.S. interpreted the cryo-EM structures. E.H., R.S., constructed and provided mutant strains. G.H. performed the evolutionary rate analysis. T.R.T., J.M.S., wrote the manuscript from the input of all other authors.

