## Supplementary Information for "The anaerobic cryo-EM structure of the methanogenic Mtr complex reveals a nitrogenase-like [Fe_8_S_9_C] cluster bound to its active site"

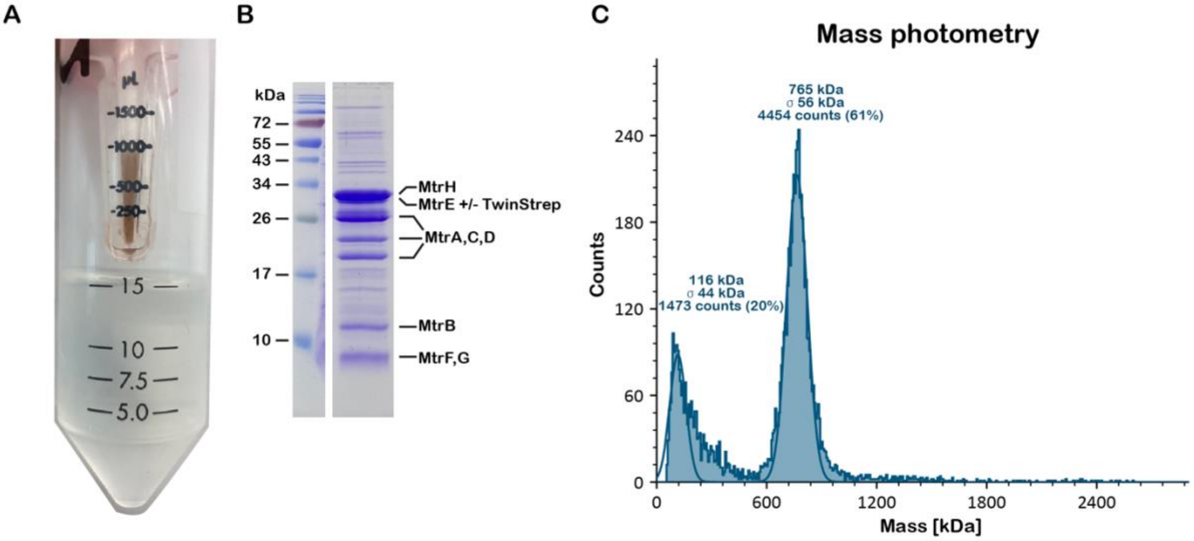

D

**ICP-MS**

**Metal concentrations**

| Sample | 59 ->59 Co [µM] | 66 ->66 Zn [µM] | 44 ->44 Ca [µM] | 56 ->56 Fe [µM] |
|---|---|---|---|---|
| Mtr H2 Anaerobic p2 | 24,69067797 | 29,79166667 | 149,2784091 | 185,5758929 |

| Sample | 60 ->60 Ni[µM] | 95 ->95 Mo[µM] | 182 ->182 W[µM] |
|---|---|---|---|
| Mtr H2 Anaerobic p2 | 1,429166667 | 0,505263158 | 0,148351648 |

**Metal/Protein ratio**

| Sample | Protein[mg/ml] | Protein[µM] | Protein[kDa] | Co / MtrABCDEFG2H | Zn / MtrABCDEFG2H |
|---|---|---|---|---|---|
| Mtr H2 Anaerobic p2 | 5 | 24,31729203 | 205615 | 1,01535475 | 1,225122708 |

| Sample | Ca / MtrABCDEFG2H | Fe / MtrABCDEFG2H | Ni / MtrABCDEFG2H | Mo / MtrABCDEFG2H | W / MtrABCDEFG2H |
|---|---|---|---|---|---|
| Mtr H2 Anaerobic p2 | 6,138776017 | 7,631437442 | 0,058771621 | 0,020777937 | 0,006100665 |

**Supplementary Fig. S1. Anaerobic purification of Mtr. (A)** 10 kDa centrifugal filter with StrepTactin eluate shows dark brown color after concentration. **(B)** 15% SDS-gel of concentrated StrepTactin eluate. **(C)** Mass photometry of anaerobically purified Mtr complex. **(D)** ICP-MS of concentrated StrepTactin eluate measured as technical triplicates. Metal/Protein ratio is calculated based on the predicted molecular weight of a MtrABCDEFGH2 protomer (205 615 Da).

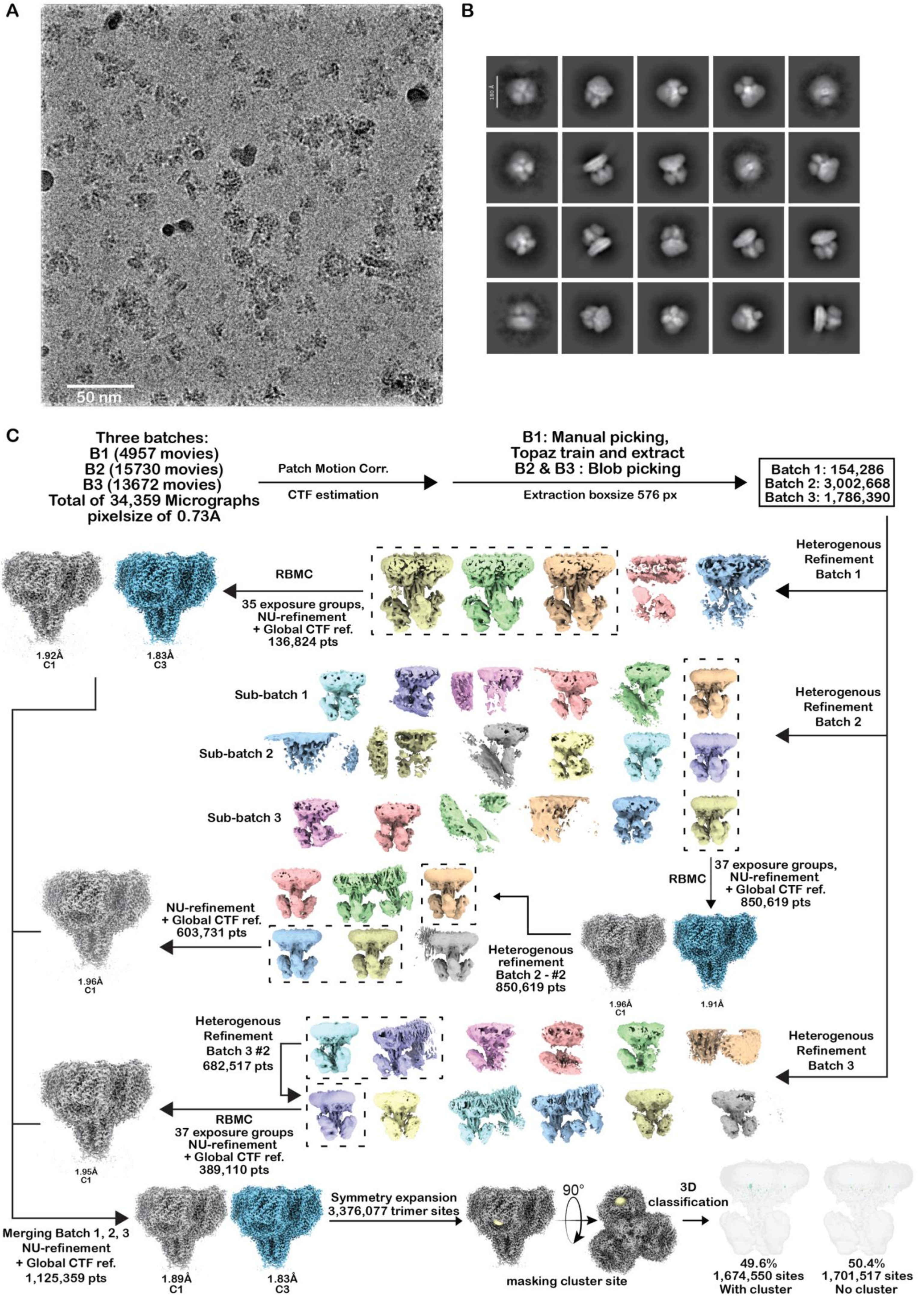
A
50 nm
B
180 Å
C
Three batches:
B1 (4957 movies)
B2 (15730 movies)
B3 (13672 movies)
Total of 34,359 Micrographs
pixelsize of 0.73Å
Patch Motion Corr.
CTF estimation
B1: Manual picking,
Topaz train and extract
B2 & B3 : Blob picking
Extraction boxsize 576 px
Batch 1: 154,286
Batch 2: 3,002,668
Batch 3: 1,786,390
Heterogenous Refinement Batch 1
RBMC
35 exposure groups,
NU-refinement
+ Global CTF ref.
136,824 pts
1.92Å
C1
1.83Å
C3
Sub-batch 1
Sub-batch 2
Sub-batch 3
Heterogenous Refinement Batch 2
37 exposure groups,
NU-refinement
+ Global CTF ref.
850,619 pts
RBMC
NU-refinement
+ Global CTF ref.
603,731 pts
1.96Å
C1
Heterogenous refinement
Batch 2 - #2
850,619 pts
1.96Å
C1
1.91Å
Heterogenous Refinement Batch 3 #2
682,517 pts
Heterogenous Refinement Batch 3
RBMC
37 exposure groups
NU-refinement
+ Global CTF ref.
389,110 pts
1.95Å
C1
Merging Batch 1, 2, 3
NU-refinement
+ Global CTF ref.
1,125,359 pts
1.89Å
C1
1.83Å
C3
Symmetry expansion
3,376,077 trimer sites
90°
masking cluster site
3D classification
49.6%
1,674,550 sites
With cluster
50.4%
1,701,517 sites
No cluster

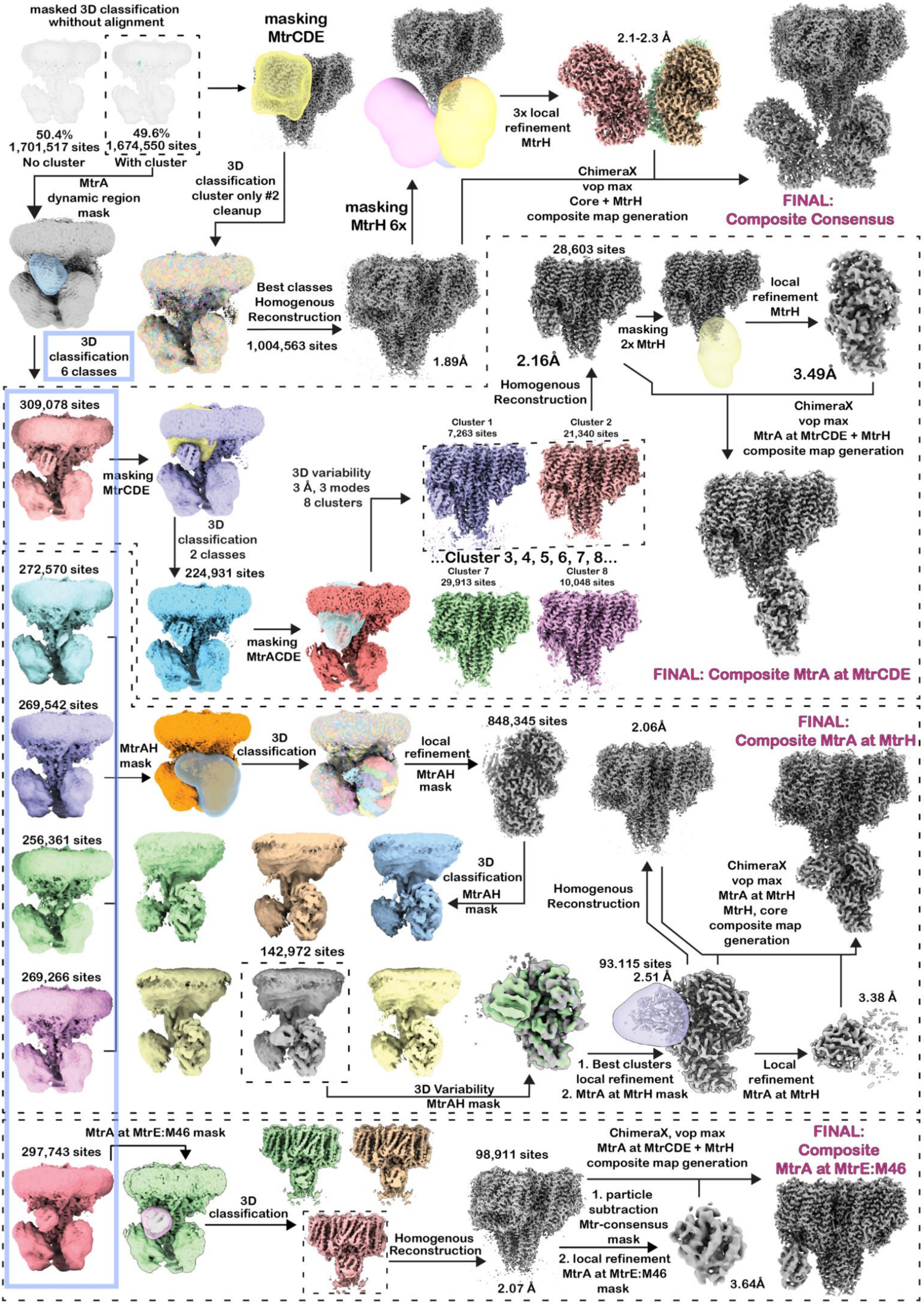

masked 3D classification whithout alignment
50.4%
1,701,517 sites
No cluster
49.6%
1,674,550 sites
With cluster
masking MtrCDE
masking MtrH 6x
3x local refinement MtrH
2.1-2.3 Å
ChimeraX vop max Core + MtrH composite map generation
FINAL: Composite Consensus
MtrA dynamic region mask
3D classification cluster only #2 cleanup
Best classes Homogenous Reconstruction
1,004,563 sites
1.89Å
28,603 sites
2.16Å
masking 2x MtrH
local refinement MtrH
3.49Å
Homogenous Reconstruction
ChimeraX vop max MtrA at MtrCDE + MtrH composite map generation
3D classification 6 classes
309,078 sites
masking MtrCDE
3D classification 2 classes
224,931 sites
3D variability 3 Å, 3 modes 8 clusters
Cluster 1 7,263 sites
Cluster 2 21,340 sites
...Cluster 3, 4, 5, 6, 7, 8...
Cluster 7 29,913 sites
Cluster 8 10,048 sites
272,570 sites
masking MtrACDE
FINAL: Composite MtrA at MtrCDE
269,542 sites
MtrAH mask
3D classification
local refinement MtrAH mask
848,345 sites
2.06Å
FINAL: Composite MtrA at MtrH
256,361 sites
3D classification MtrAH mask
Homogenous Reconstruction
ChimeraX vop max MtrA at MtrH MtrH, core composite map generation
269,266 sites
142,972 sites
93.115 sites
2.51 Å
3.38 Å
3D Variability MtrAH mask
1. Best clusters local refinement 2. MtrA at MtrH mask
Local refinement MtrA at MtrH
MtrA at MtrE:M46 mask
297,743 sites
3D classification
Homogenous Reconstruction
98,911 sites
2.07 Å
ChimeraX, vop max MtrA at MtrCDE + MtrH composite map generation
1. particle subtraction Mtr-consensus mask
2. local refinement MtrA at MtrE:M46 mask
3.64Å
FINAL: Composite MtrA at MtrE:M46

**Supplementary Fig. S2. CryoEM data processing workflow. (A)** Exemplary micrograph with 50 nm scale bar. **(B)** Selected 2D class averages showing different particle orientations. **(C)** Mtr data processing tree.

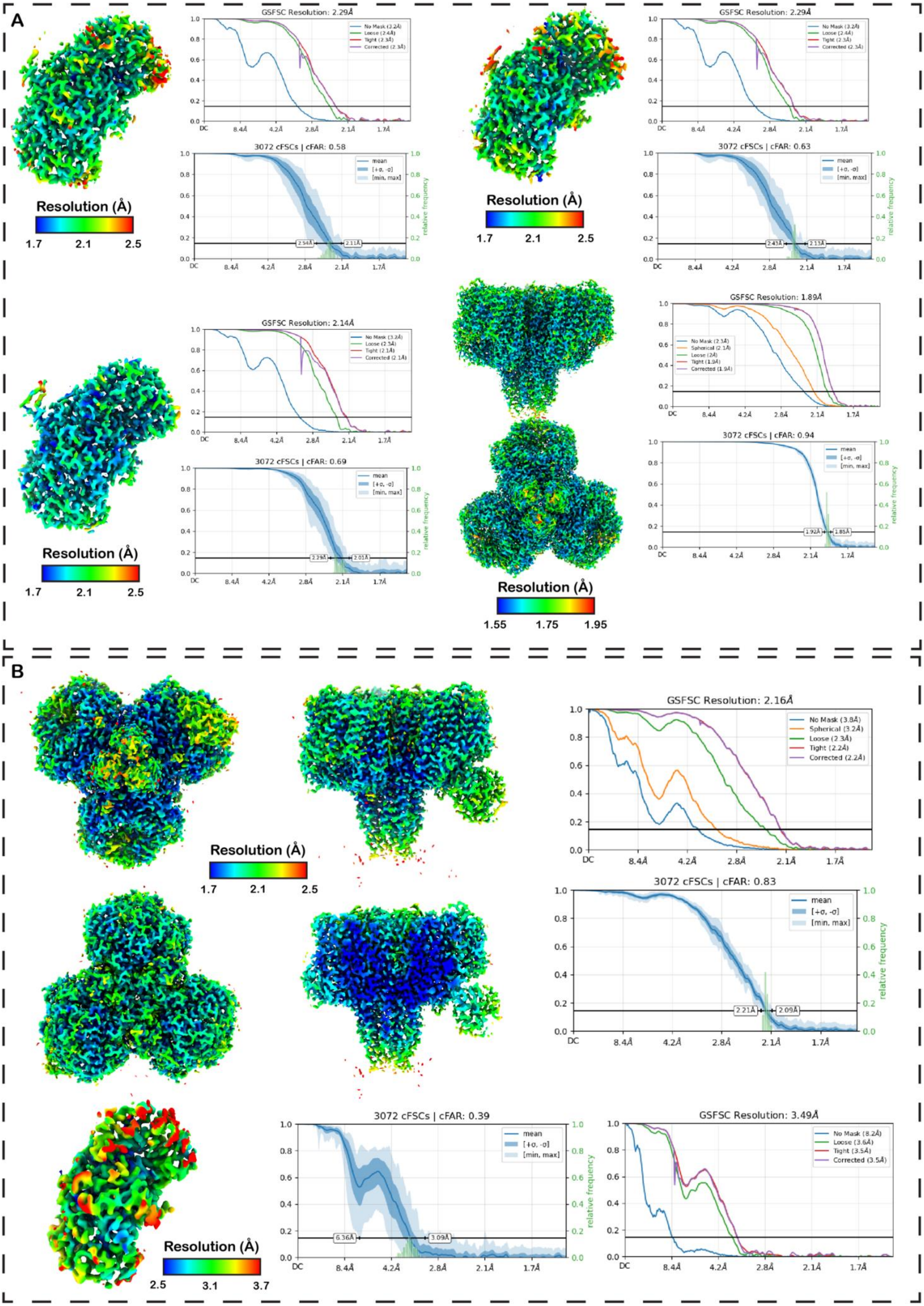

A
GSFSC Resolution: 2.29Å
No Mask (3.2Å)
Loose (2.4Å)
Tight (2.3Å)
Corrected (2.3Å)
3072 cFSCs | cFAR: 0.58
mean
[+σ, -σ]
[min, max]
relative frequency
2.54Å
2.11Å
Resolution (Å)
1.7
2.1
2.5
GSFSC Resolution: 2.29Å
3072 cFSCs | cFAR: 0.63
2.43Å
2.13Å
GSFSC Resolution: 2.14Å
No Mask (3.2Å)
Loose (2.3Å)
Tight (2.1Å)
Corrected (2.1Å)
3072 cFSCs | cFAR: 0.69
2.29Å
2.01Å
GSFSC Resolution: 1.89Å
No Mask (2.3Å)
Spherical (2.1Å)
Loose (2Å)
Tight (1.9Å)
Corrected (1.9Å)
3072 cFSCs | cFAR: 0.94
1.92Å
1.85Å
1.55
1.75
1.95
DC
8.4Å
4.2Å
2.8Å
2.1Å
1.7Å
B
GSFSC Resolution: 2.16Å
No Mask (3.8Å)
Spherical (3.2Å)
Loose (2.3Å)
Tight (2.2Å)
Corrected (2.2Å)
3072 cFSCs | cFAR: 0.83
2.21Å
2.09Å
3072 cFSCs | cFAR: 0.39
6.36Å
3.09Å
GSFSC Resolution: 3.49Å
No Mask (8.2Å)
Loose (3.6Å)
Tight (3.5Å)
Corrected (3.5Å)
2.5
3.1
3.7

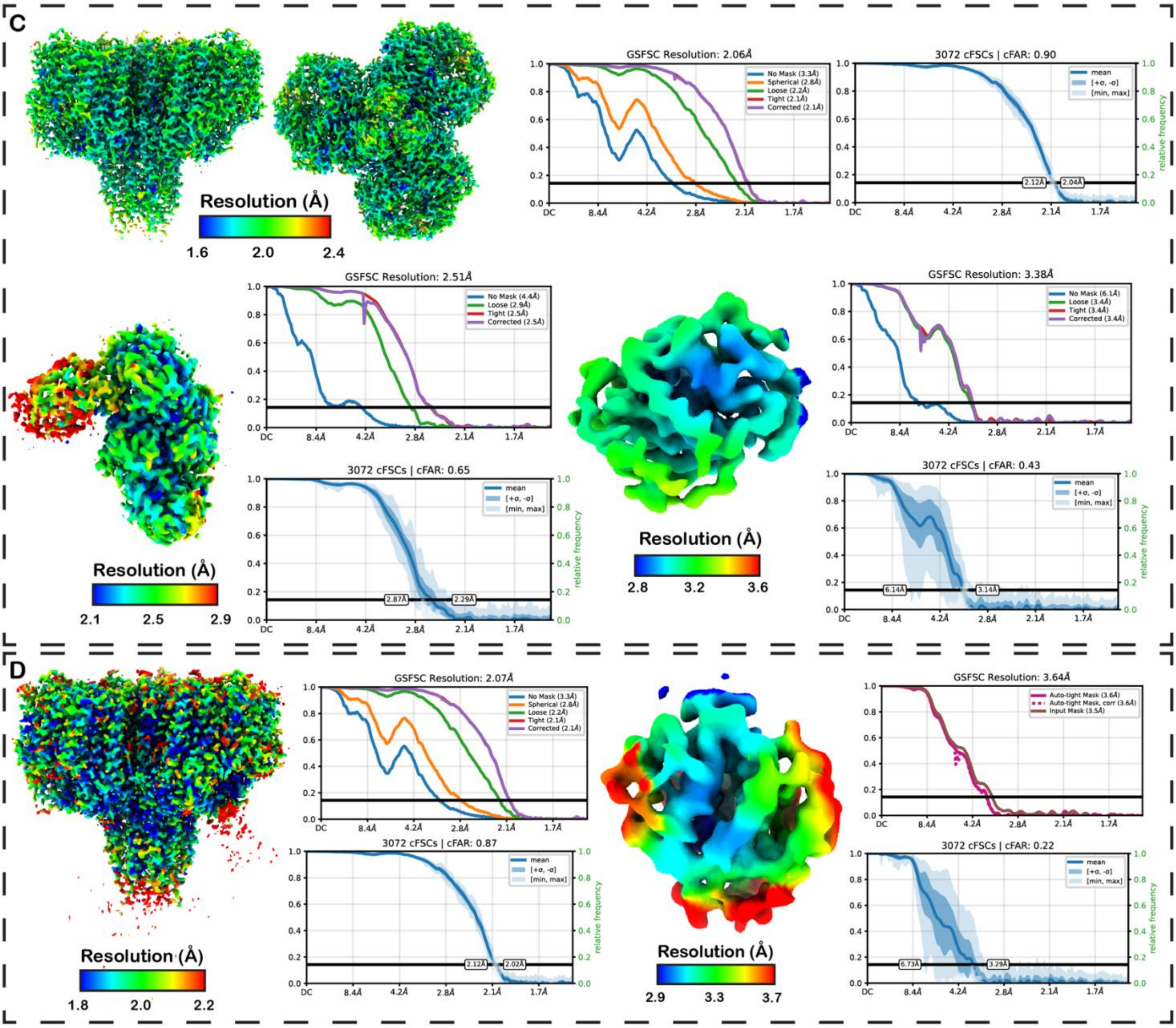

**Supplementary Fig. S3. Cryo-EM maps resolution and orientation diagnostics. (A)** Consensus: Mtr core map (different views), MtrH maps. **(B)** MtrA at MtrCDE: Mtr core map (different views and cutaway view), MtrH maps **(C)** MtrA at MtrH: Mtr core map (different views), MtrH map, MtrA at MtrH map **(D)** MtrA at MtrE:M46: Mtr core map, MtrA at MtrE:M46 map.

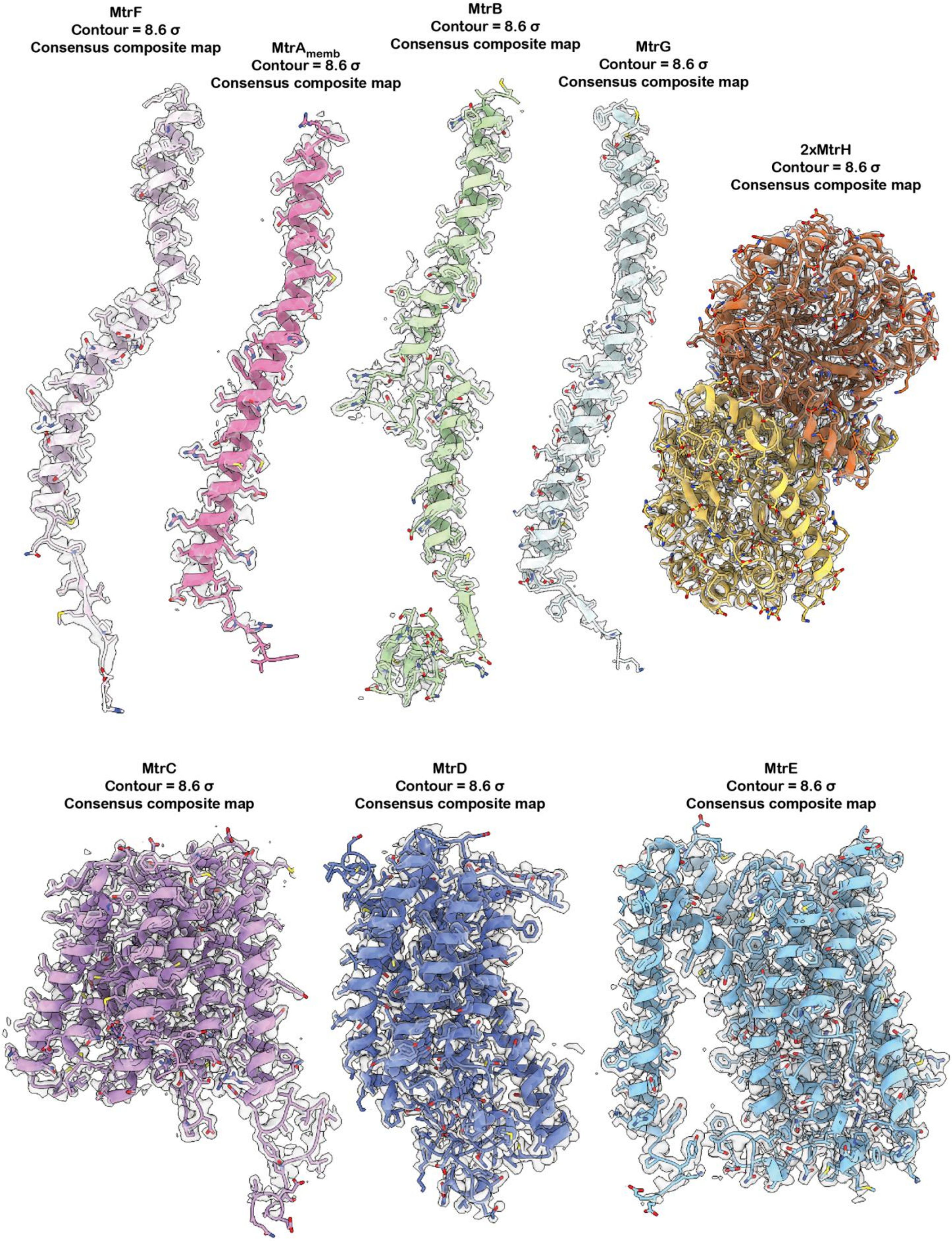
MtrF
Contour = 8.6 σ
Consensus composite map
MtrA$_{memb}$
Contour = 8.6 σ
Consensus composite map
MtrB
Contour = 8.6 σ
Consensus composite map
MtrG
Contour = 8.6 σ
Consensus composite map
2xMtrH
Contour = 8.6 σ
Consensus composite map
MtrC
Contour = 8.6 σ
Consensus composite map
MtrD
Contour = 8.6 σ
Consensus composite map
MtrE
Contour = 8.6 σ
Consensus composite map

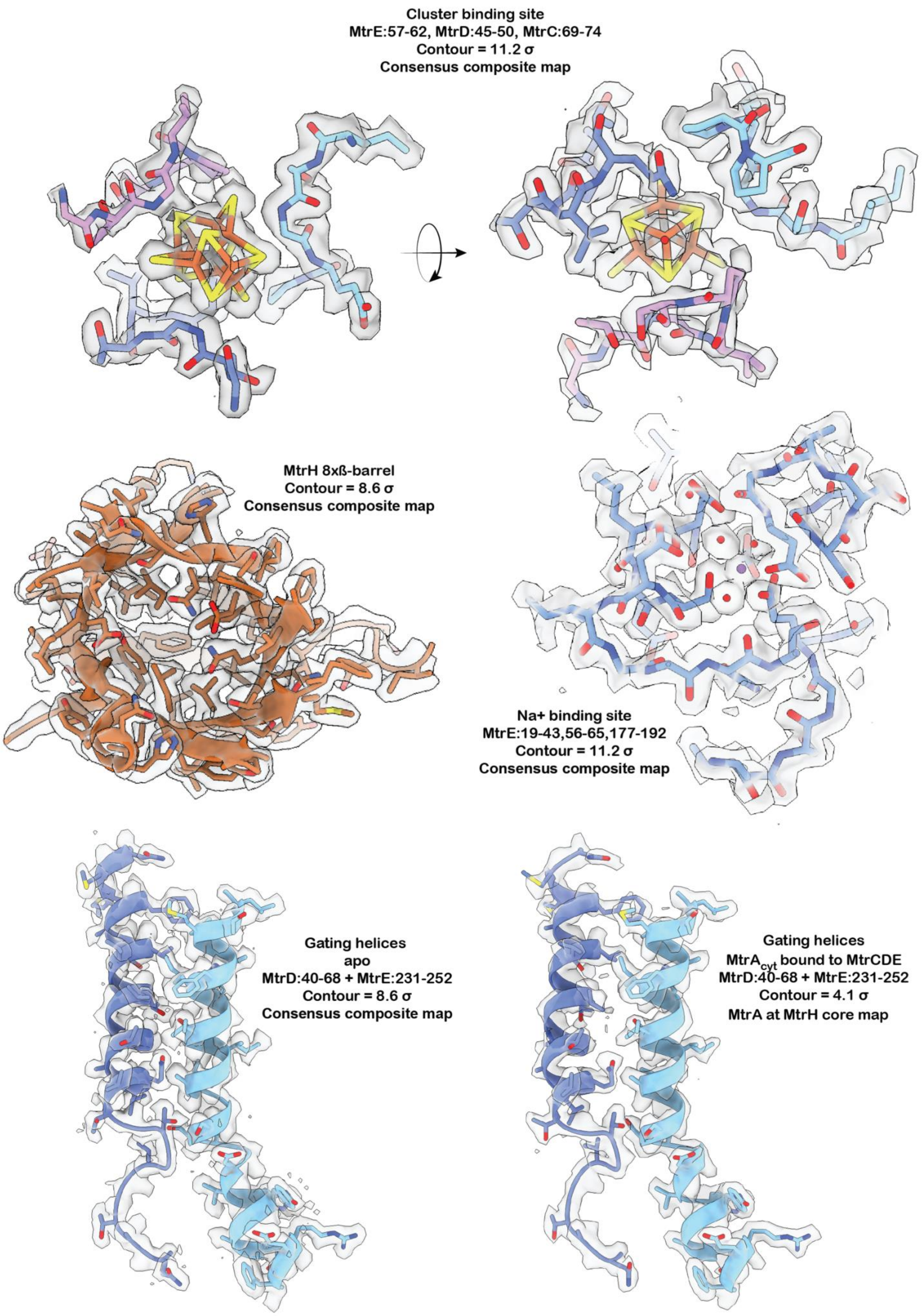
Cluster binding site
MtrE:57-62, MtrD:45-50, MtrC:69-74
Contour = 11.2 σ
Consensus composite map
MtrH 8xß-barrel
Contour = 8.6 σ
Consensus composite map
Na+ binding site
MtrE:19-43,56-65,177-192
Contour = 11.2 σ
Consensus composite map
Gating helices
apo
MtrD:40-68 + MtrE:231-252
Contour = 8.6 σ
Consensus composite map
Gating helices
MtrA$_{cyt}$ bound to MtrCDE
MtrD:40-68 + MtrE:231-252
Contour = 4.1 σ
MtrA at MtrH core map

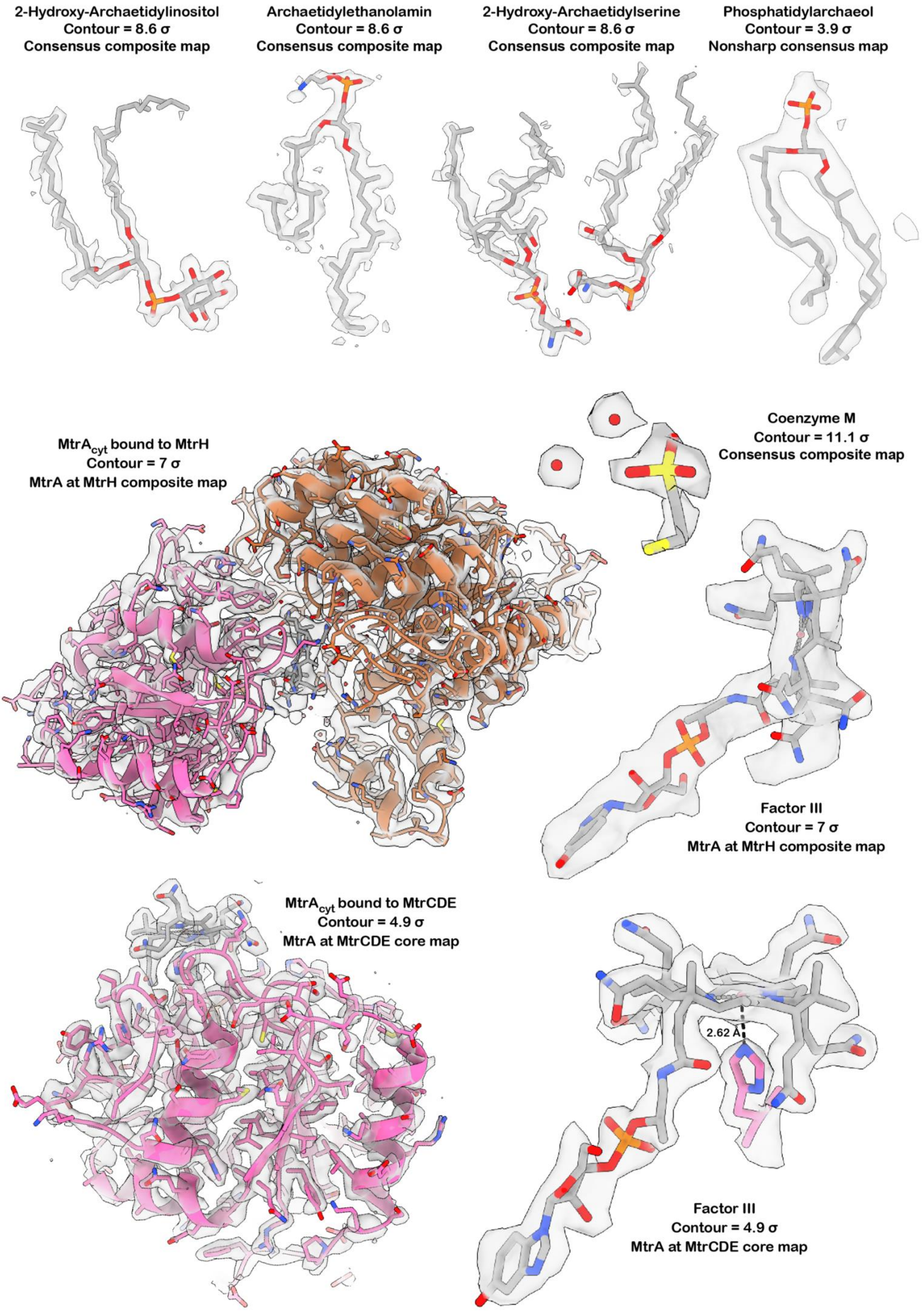
2-Hydroxy-Archaetidylinositol
Contour = 8.6 σ
Consensus composite map
Archaetidylethanolamin
Contour = 8.6 σ
Consensus composite map
2-Hydroxy-Archaetidylserine
Contour = 8.6 σ
Consensus composite map
Phosphatidylarchaeol
Contour = 3.9 σ
Nonsharp consensus map
MtrA$_{cyt}$ bound to MtrH
Contour = 7 σ
MtrA at MtrH composite map
Coenzyme M
Contour = 11.1 σ
Consensus composite map
Factor III
Contour = 7 σ
MtrA at MtrH composite map
MtrA$_{cyt}$ bound to MtrCDE
Contour = 4.9 σ
MtrA at MtrCDE core map
2.62 Å
Factor III
Contour = 4.9 σ
MtrA at MtrCDE core map

**Supplementary Fig. S4. Model-to-map fit.** Cryo-EM density maps surrounding the Mtr subunits, associated cofactors, and selected regions are shown at the indicated sigma thresholds used for visualization. These representations illustrate the quality of the respective reconstructions.

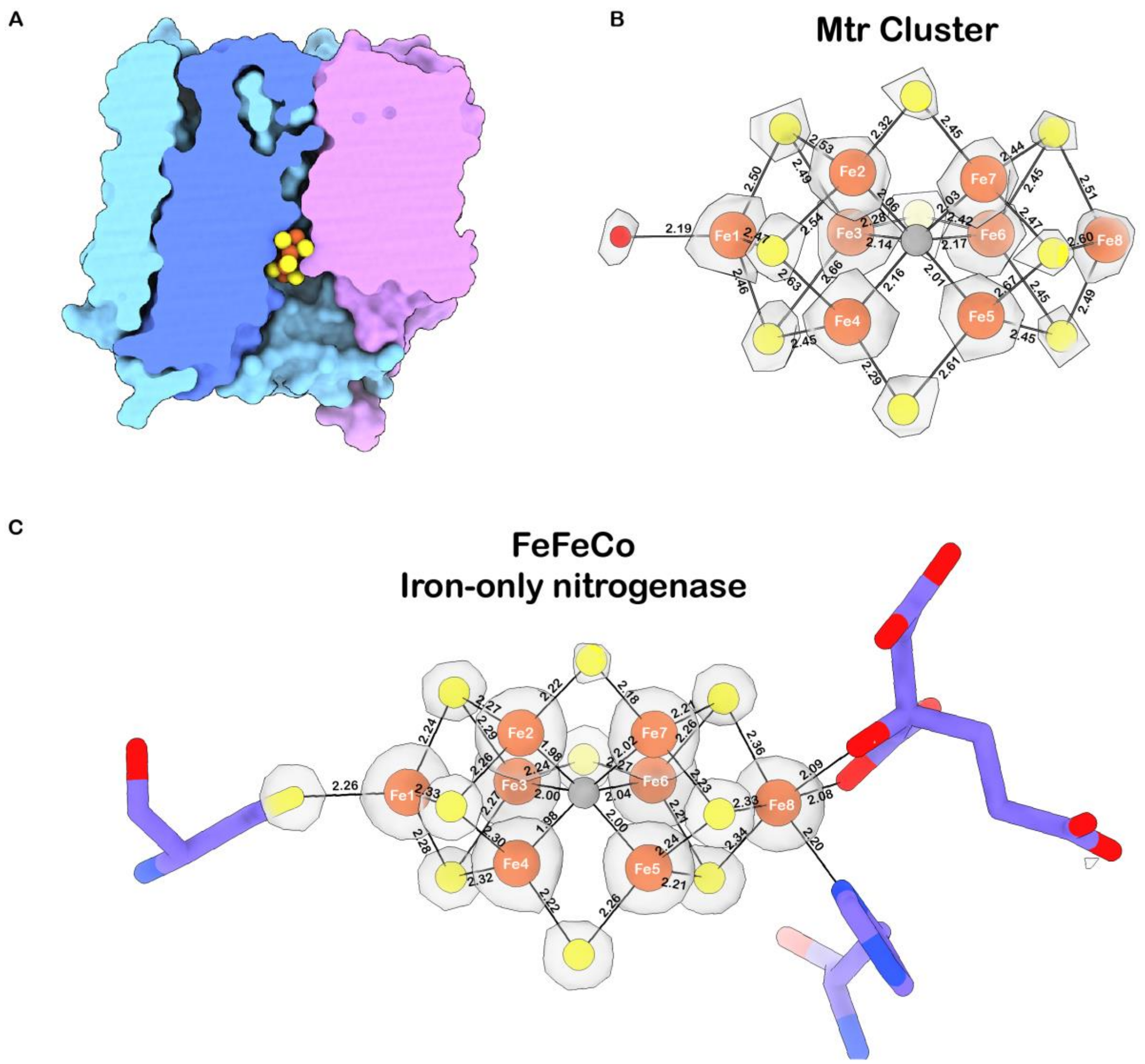

**Supplementary Fig. S5. Structural details of the metallocluster (A)** Cutaway view of the MtrCDE trimer in surface representation, revealing embedding of the metallocluster within the trimer. **(B)** High-threshold cryo-em density (contour level = 30.4 σ) together with the atomic model of the [$Fe_8S_9C$]-L-cluster, including a water ligand (red) coordinated to the apical iron. Bond distances are indicated in Å. **(C)** 2mFo–DFc electron density map of the FeFe Cofactor from the iron-only nitrogenase together with the atomic model (PDB: 8BOQ) of the metal centers, coordinating protein ligands and homocitrate. Bond distances are indicated in Å. The Fe–S bond distances in the Mtr cluster are 5–15% longer than those in FeFeCo.

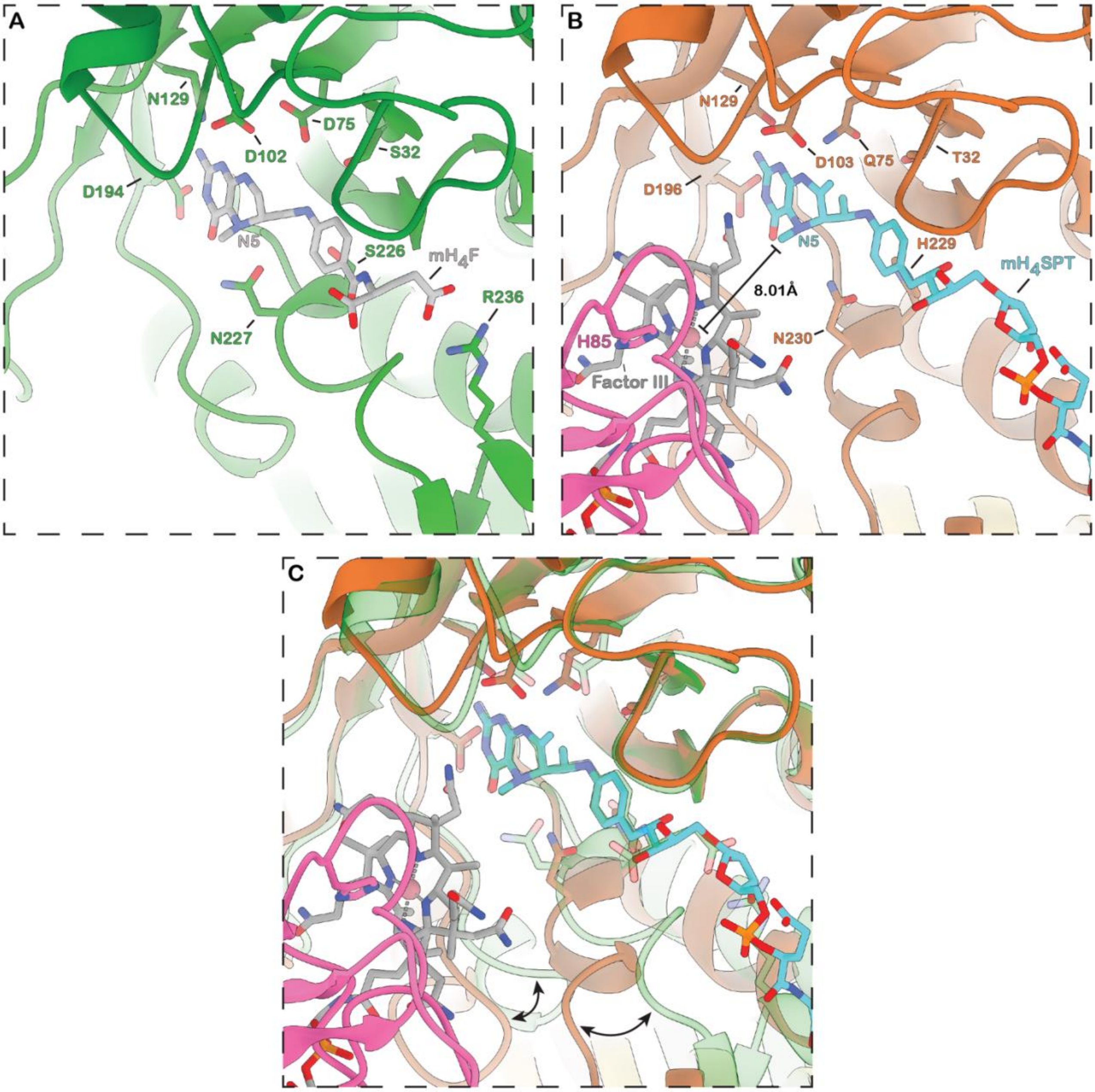

**Supplementary Fig. S6. Active site rearrangements in MtrH inferred from comparison with MtgA (A)** Close-up view of the active site of MtgA (PDB: 6SJN) with methyl-tetrahydrofolate ($mH_4F$) bound and catalytically important residues highlighted. **(B)** Close-up view of the MtrH active site in the MtrA-MtrH structure, with methyl-tetrahydrosarcinapterin ($mH_4SPT$) modelled based on the electron density of $mH_4F$ bound to MtgA. The distance between the corrinoid cobalt and the reactive methyl group is indicated and catalytically important residues are highlighted. **(C)** Superposition of the active site of MtgA and MtrH indicates that substrate-binding elements (MtrH:197-205 and MtrH:228-260) in $mH_4SPT$-free MtrH must undergo rearrangements to enable proper substrate binding. These changes likely allow deeper insertion of the corrinoid into the TIM barrel, thereby shortening the cobalt–methyl distance, and facilitating methyl transfer.

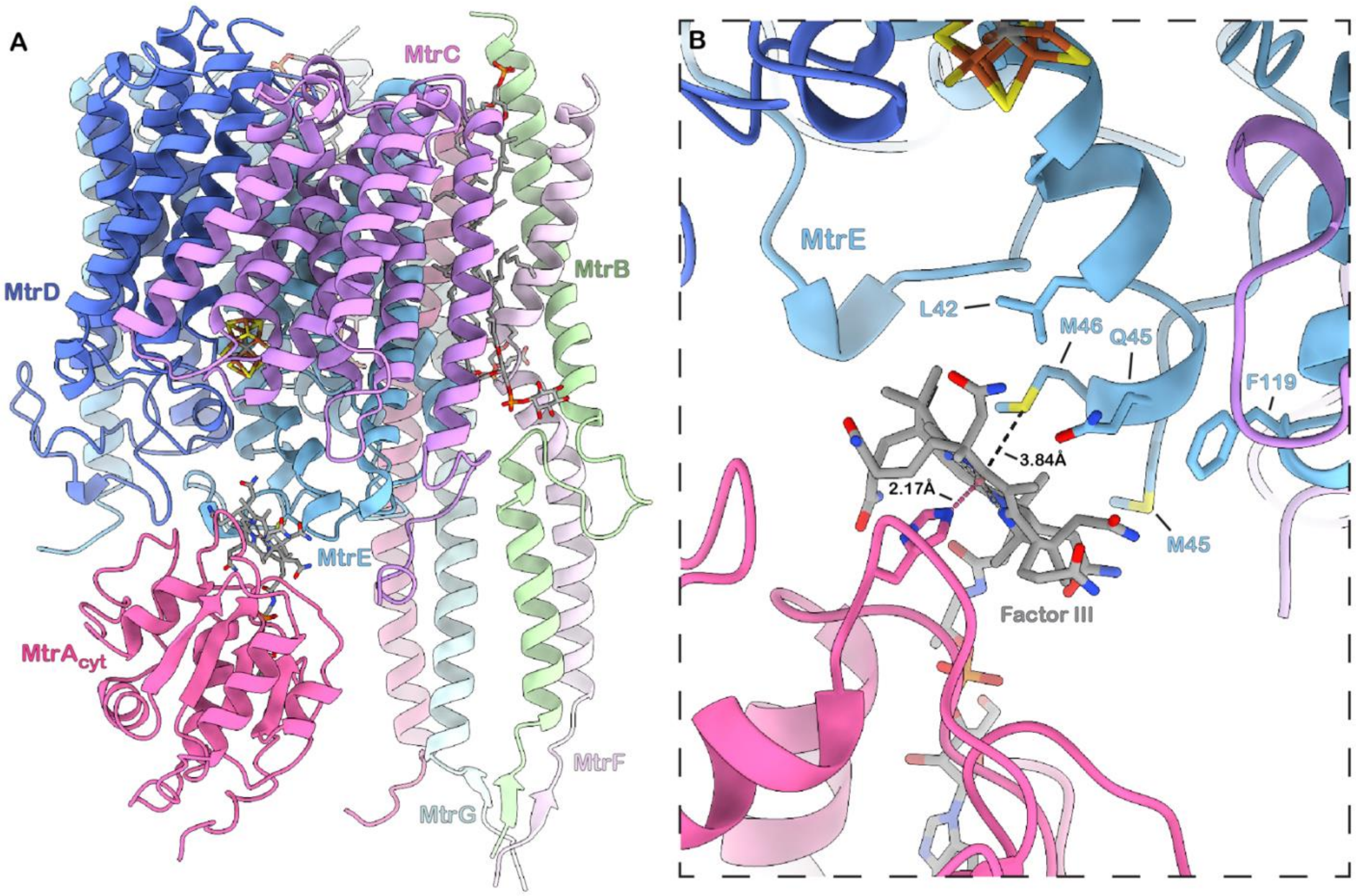

**Supplementary Fig. S7. interaction between the MtrA corrinoid and MtrE:M46. (A)** Side view of a single protomer within the trimeric Mtr complex showing MtrA positioned adjacent to MtrE:M46 at the cytosolic helix. **(B)** Close-up of the MtrA-MtrE:M46 interaction. The cobalt-sulfur distance distance of 3.84 Å is inconsistent with a direct coordination of methionine to the corrinoid but the interaction appears to be mediated by a combination of nonpolar und polar contacts between the corrinoid and residues L42, M45, M46, and Q45.

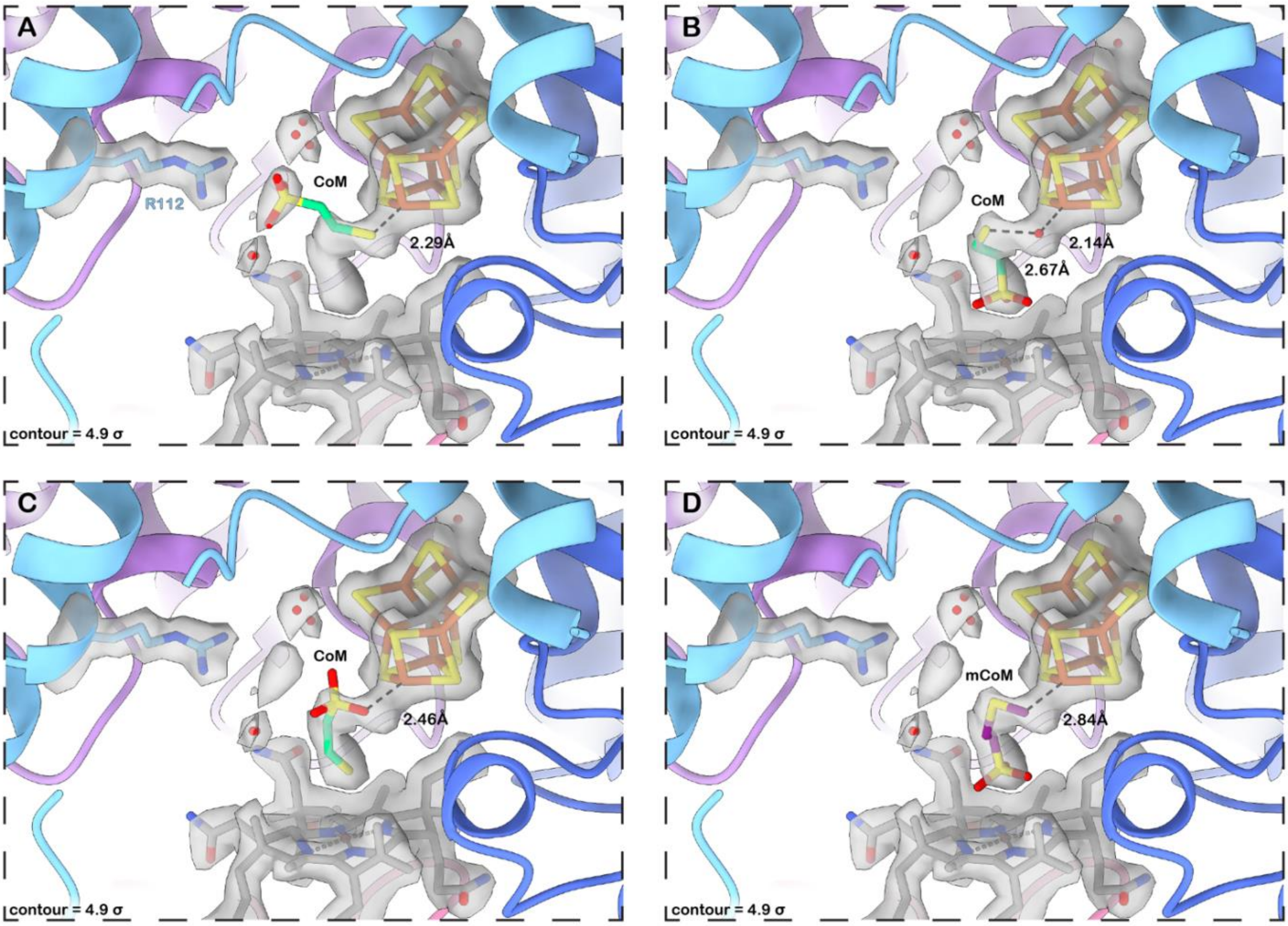

**Supplementary Fig. S8. Potential CoM/mCoM binding modes consistent with the ambiguous density at the apical-iron. (A)** CoM interacts with its sulfonate group with the guanidinium head-group of Arg112, while its sulfhydryl is oriented toward the apical iron. **(B)** In an alternative orientation, the sulfonate group is directed toward the corrinoid cobalt, while the sulfhydryl points toward the apical iron, potentially bridged by an additional ligand (e.g. water). **(C)** A further possibility places the sulfonate toward the apical iron and the sulfhydryl toward the cobalt-corrinoid. **(D)** Alternatively, the observed density may correspond to methyl-CoM formed after methyl transfer, with the methyl-group oriented toward the apical iron and the sulfonate pointing toward the corrinoid cobalt.

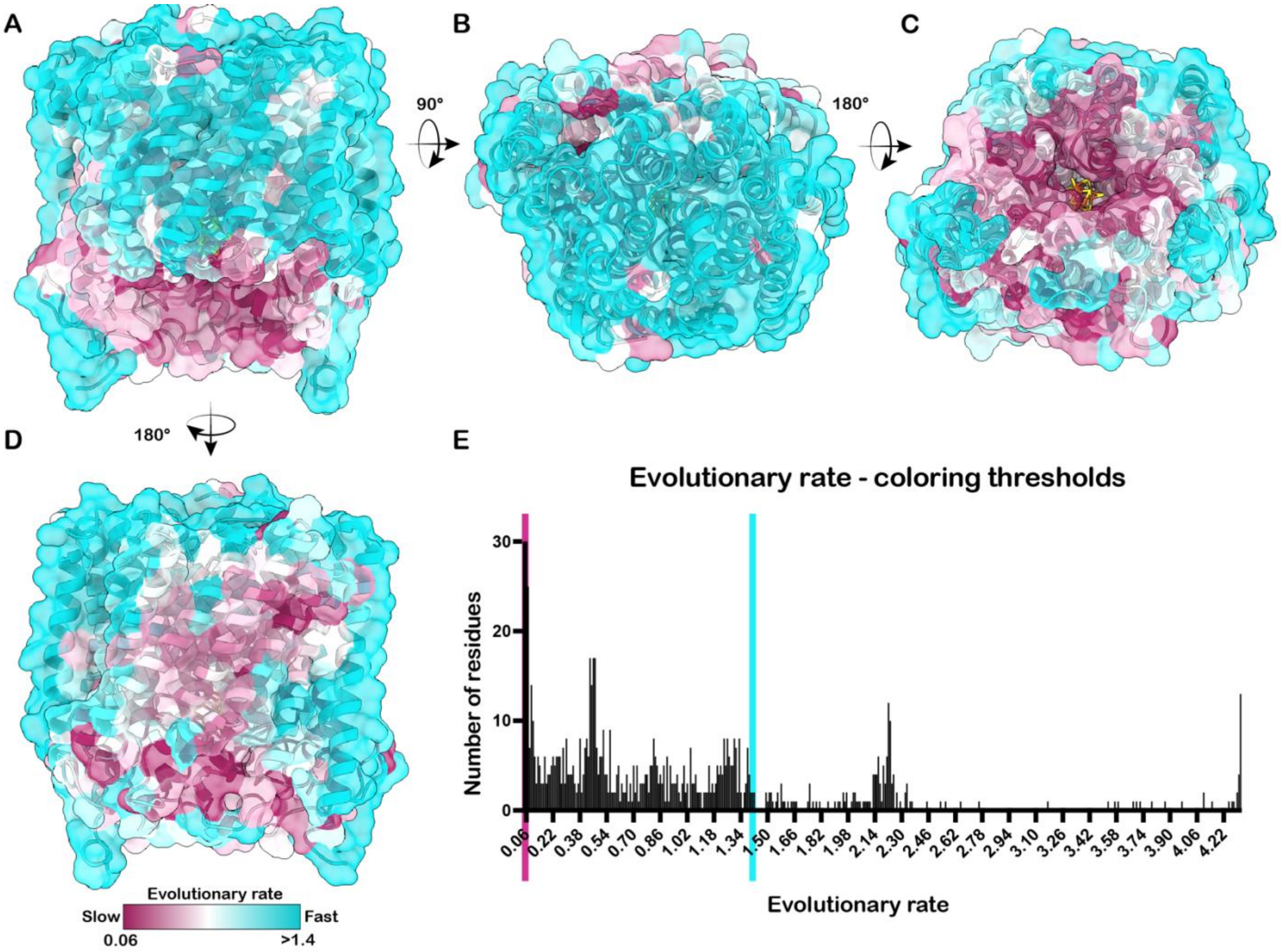

**Supplementary Fig. S9. Evolutionary rate mapping on the MtrCDE reveals surface conservation of the trimer. (A)** Side view showing that the membrane facing regions of MtrC and MtrD and **(B)** the extracellular site of the trimer are fast-evolving. **(C)** MtrA and metallocluster binding site at the cytosolic region shows strong conservation. **(D)** Stalk-facing surface of the trimer mainly formed by MtrE contains both slow and fast evolving residues. **(E)** Histogram showing the distribution of residues across evolutionary rates. The lower and upper thresholds used for color mapping are indicated by pink (0.06) and cyan (1.4) vertical lines.

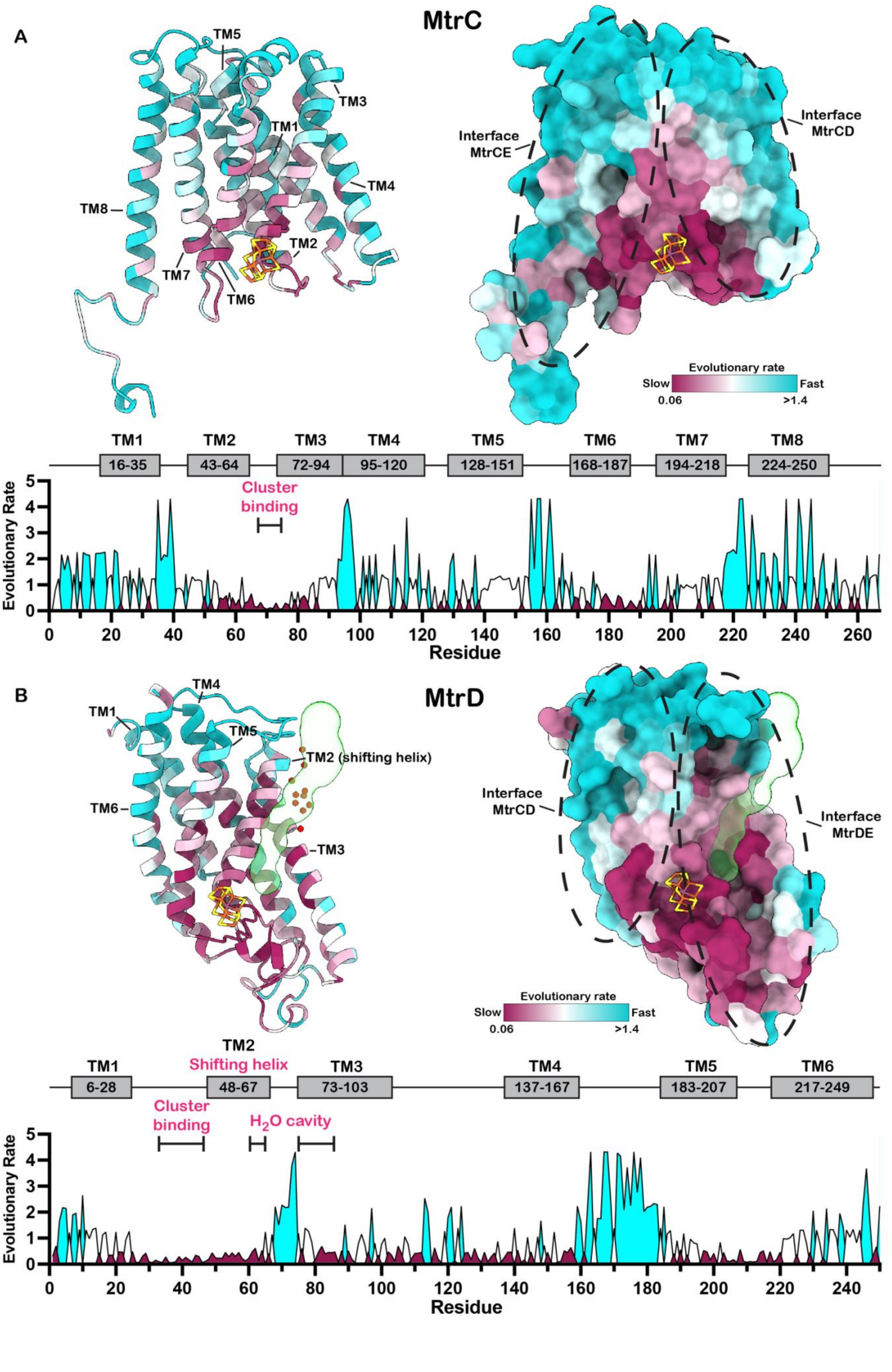
A
MtrC
TM5
TM3
TM1
TM4
TM8
TM2
TM7
TM6
Interface MtrCE
Interface MtrCD
Evolutionary rate
Slow
Fast
0.06
>1.4
TM1 16-35
TM2 43-64
TM3 72-94
TM4 95-120
TM5 128-151
TM6 168-187
TM7 194-218
TM8 224-250
Cluster binding
Evolutionary Rate
Residue
B
MtrD
TM4
TM1
TM5
TM2 (shifting helix)
TM6
TM3
Interface MtrCD
Interface MtrDE
Evolutionary rate
Slow
Fast
0.06
>1.4
TM1 6-28
TM2 Shifting helix 48-67
TM3 73-103
TM4 137-167
TM5 183-207
TM6 217-249
Cluster binding
H2O cavity
Evolutionary Rate
Residue

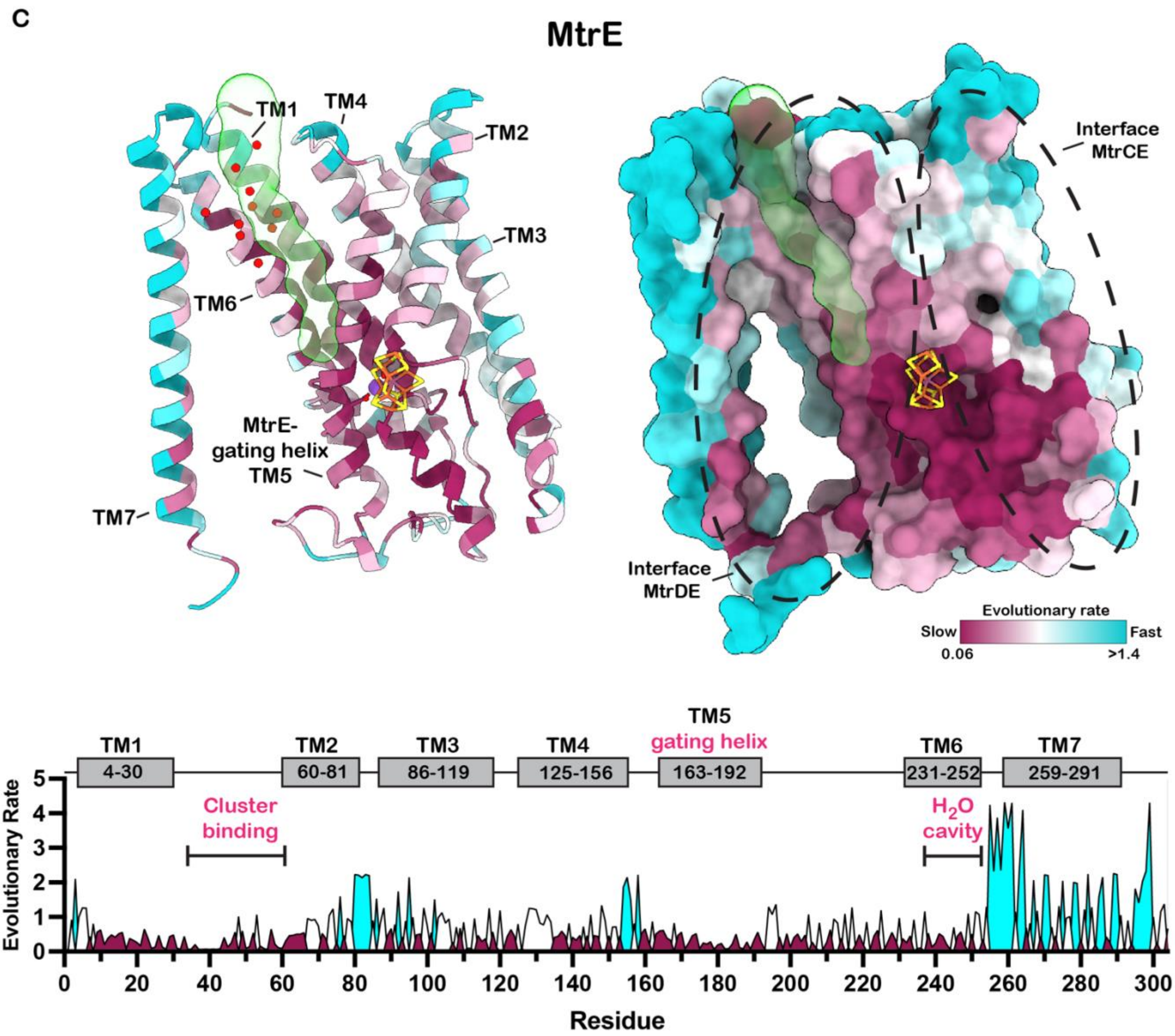

**Supplementary Fig. S10. Subunit-specific evolutionary conservation in the MtrA–MtrCDE complex. (A) MtrC. Left:** Cartoon representation with transmembrane helices (TM1–8) labeled and the metallocluster shown. **Right:** Surface representation with MtrCE and MtrCD interface regions indicated (dotted ellipsoids). **Bottom:** Evolutionary rate profile of all residues. Evolutionary rates <0.7 are colored maroon, 0.7–1.4 white, and >1.4 cyan, with a 1D helix map and selected regions highlighted. **(B) MtrD. Left:** Cartoon representation with transmembrane helices (TM1–6), metallocluster, and predicted tunnel (transparent green). Waters at the extracellular cavity/tunnel exit are shown. **Right:** Surface representation with MtrCD and MtrDE interfaces indicated and tunnel shown. **Bottom**: Evolutionary rate profile as in (A). **(C) MtrE. Left:** Cartoon representation with transmembrane helices (TM1–7), metallocluster, sodium ion, and predicted tunnel (transparent green). Waters at the extracellular cavity/tunnel exit are shown. **Right:** Surface representation with MtrCE and MtrDE interfaces indicated and tunnel shown. **Bottom:** Evolutionary rate profile as in (A).

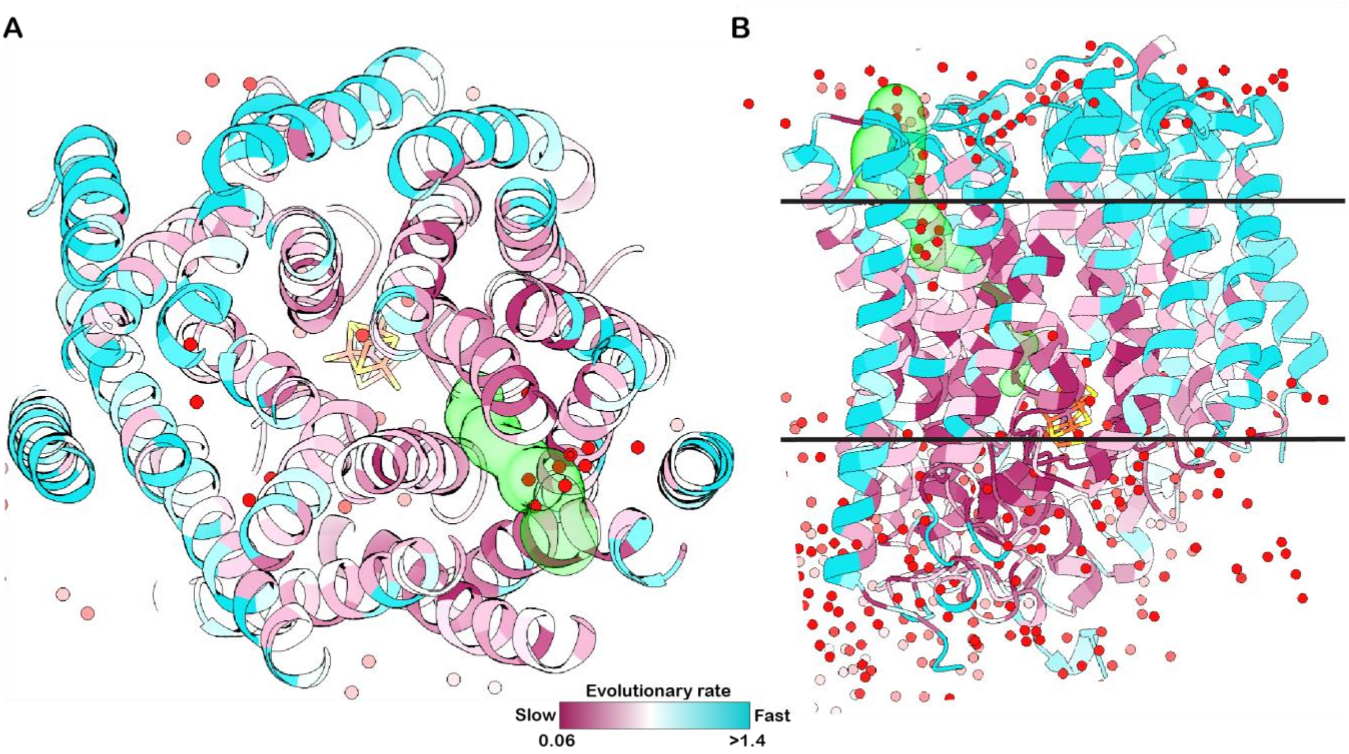

**Supplementary Fig. S11. Ion-channel between MtrD and MtrE supported by evolutionary rate mapping. (A)** Top view from the extracellular of the MtrCDE trimer, highlighting strong conservation at the MtrCD interface, particularly around the extracellular water cavity and the predicted tunnel. In contrast, other regions, including the MtrCE and MtrCD interfaces, are less conserved, suggesting they are less likely to function as ion channel pathways. **(B)** Side view of the trimer with clipping planes used in (A) indicated as horizontal lines

### Cryo-EM data collection, refinement and validation statistics

| A | **Mtr apo Composite map** (EMD-58097) (PDB: 30VS) | **Mtr apo consensus map** (EMD-58079) | **MtrH local map #1** (EMD-58094) | **MtrH local map #2** (EMD-58081) | **MtrH local map #3** (EMDB-58084) |
|---|---|---|---|---|---|
| **Data collection and processing** | | | | | |
| Magnification | 165 000x | 165 000x | 165 000x | 165 000x | 165 000x |
| Voltage (kV) | 300 | 300 | 300 | 300 | 300 |
| Electron exposure (e–/Å$^2$) | 55 | 55 | 55 | 55 | 55 |
| Defocus range (µm) | 0.5-3.5 | 0.5-3.5 | 0.5-3.5 | 0.5-3.5 | 0.5-3.5 |
| Pixel size (Å) | 0.73 | 0.73 | 0.73 | 0.73 | 0.73 |
| Symmetry imposed | C1 | C1 | C1 | C1 | C1 |
| Initial particle images (no.) | 3 376 077 | 3 376 077 | 3 376 077 | 3 376 077 | 3 376 077 |
| Final particle images (no.) | 1 004 563 | 1 004 563 | 1 004 563 | 1 004 563 | 1 004 563 |
| Map resolution (Å) | | 1.89 | 2.14 | 2.29 | 2.29 |
| FSC threshold | | 0.143 | 0.143 | 0.143 | 0.143 |
| Map resolution range (Å) | | ~1.55-2.0 | ~1.7-2.3 | ~1.8-2.5 | ~1.8-2.5 |
| **Refinement** | | | | | |
| Initial model (PDB code) | 9QTS | | | | |
| Model resolution (Å) | | | | | |
| FSC threshold | | | | | |
| Model resolution range (Å) | | | | | |
| Map sharpening *B* factor (Å$^2$) | | 49.6 | 62.3 | 62.1 | 64.1 |
| Model composition | | | | | |
| Non-hydrogen atoms | 42052 | | | | |
| Protein residues | 5217 | | | | |
| Ligands | 24 | | | | |
| *B* factors (Å$^2$) | | | | | |
| Protein | 47.45 | | | | |
| Ligand | 47.01 | | | | |
| R.m.s. deviations | | | | | |
| Bond lengths (Å) | 0.002 | | | | |
| Bond angles (°) | 0.489 | | | | |
| Validation | | | | | |
| MolProbity score | 1.73 | | | | |
| Clashscore | 12.59 | | | | |
| Poor rotamers (%) | 0.73 | | | | |
| Ramachandran plot | | | | | |
| Favored (%) | 97.37 | | | | |
| Allowed (%) | 2.58 | | | | |
| Disallowed (%) | 0.06 | | | | |

### Cryo-EM data collection, refinement and validation statistics

| | **MtrA at MtrCDE Composite map** (EMD-58098) (PDB: 30VT) | **MtrA at MtrCDE consensus map** (EMD-58092) | **MtrH local map** (EMD-58095) |
|---|---|---|---|
| **Data collection and processing** | | | |
| Magnification | 165 000x | 165 000x | 165 000x |
| Voltage (kV) | 300 | 300 | 300 |
| Electron exposure (e–/Å$^2$) | 55 | 55 | 55 |
| Defocus range (μm) | 0.5-3.5 | 0.5-3.5 | 0.5-3.5 |
| Pixel size (Å) | 0.73 | 0.73 | 0.73 |
| Symmetry imposed | C1 | C1 | C1 |
| Initial particle images (no.) | 3 376 077 | 3 376 077 | 3 376 077 |
| Final particle images (no.) | 28 603 | 28 603 | 28 603 |
| Map resolution (Å) | | 2.16 | 3.49 |
| FSC threshold | | 0.143 | 0.143 |
| Map resolution range (Å) | | ~1.7-2.3 | ~2.6-3.7 |
| **Refinement** | | | |
| Initial model (PDB code) | 9QTS | | |
| Model resolution (Å) | | | |
| FSC threshold | | | |
| Model resolution range (Å) | | | |
| Map sharpening *B* factor (Å$^2$) | | 30.8 | 72.3 |
| Model composition | | | |
| Non-hydrogen atoms | 32141 | | |
| Protein residues | 4056 | | |
| Ligands | 25 | | |
| *B* factors (Å$^2$) | | | |
| Protein | 93.73 | | |
| Ligand | 75.27 | | |
| R.m.s. deviations | | | |
| Bond lengths (Å) | 0.006 | | |
| Bond angles (°) | 0.700 | | |
| Validation | | | |
| MolProbity score | 2.09 | | |
| Clashscore | 15.21 | | |
| Poor rotamers (%) | 2.77 | | |
| Ramachandran plot | | | |
| Favored (%) | 97.81 | | |
| Allowed (%) | 1.73 | | |
| Disallowed (%) | 0.00 | | |

### Cryo-EM data collection, refinement and validation statistics

| | **MtrA at MtrH Composite map** (EMD-58111) (PDB: 30WF) | **MtrA at MtrH consensus map** (EMD-58078) | **MtrH local map** (EMD-58107) | **MtrA local map** (EMD-58080) |
|---|---|---|---|---|
| **Data collection and processing** | | | | |
| Magnification | 165 000x | 165 000x | 165 000x | 165 000x |
| Voltage (kV) | 300 | 300 | 300 | 300 |
| Electron exposure (e–/Å$^2$) | 55 | 55 | 55 | 55 |
| Defocus range (µm) | 0.5-3.5 | 0.5-3.5 | 0.5-3.5 | 0.5-3.5 |
| Pixel size (Å) | 0.73 | 0.73 | 0.73 | 0.73 |
| Symmetry imposed | C1 | C1 | C1 | C1 |
| Initial particle images (no.) | 3 376 077 | 3 376 077 | 3 376 077 | 3 376 077 |
| Final particle images (no.) | 93 115 | 93 115 | 93 115 | 93 115 |
| Map resolution (Å) | | 2.06 | 2.51 | 3.38 |
| FSC threshold | | 0.143 | 0.143 | 0.143 |
| Map resolution range (Å) | | ~1.7-2.2 | ~2.1-2.9 | ~2.8-3.3 |
| **Refinement** | | | | |
| Initial model (PDB code) | 9QTS | | | |
| Model resolution (Å) | | | | |
| FSC threshold | | | | |
| Model resolution range (Å) | | | | |
| Map sharpening *B* factor (Å$^2$) | | 58.8 | 62.2 | 80 |
| Model composition | | | | |
| Non-hydrogen atoms | 32595 | | | |
| Protein residues | 4064 | | | |
| Ligands | 25 | | | |
| *B* factors (Å$^2$) | | | | |
| Protein | 124.91 | | | |
| Ligand | 118.98 | | | |
| R.m.s. deviations | | | | |
| Bond lengths (Å) | 0.002 | | | |
| Bond angles (°) | 0.506 | | | |
| Validation | | | | |
| MolProbity score | 1.83 | | | |
| Clashscore | 16.49 | | | |
| Poor rotamers (%) | 1.24 | | | |
| Ramachandran plot | | | | |
| Favored (%) | 97.83 | | | |
| Allowed (%) | 2.14 | | | |
| Disallowed (%) | 0.02 | | | |

### Cryo-EM data collection, refinement and validation statistics

| | **MtrA at MtrE:M46 Composite map** (EMD-58076) (PDB: 30UY) | **MtrA at MtrE:M46 consensus map** (EMD-58074) | **MtrA at MtrE:M46 Local map** (EMD-58075) |
|---|---|---|---|
| **Data collection and processing** | | | |
| Magnification | 165 000x | 165 000x | 165 000x |
| Voltage (kV) | 300 | 300 | 300 |
| Electron exposure (e–/Å$^2$) | 55 | 55 | 55 |
| Defocus range (μm) | 0.5-3.5 | 0.5-3.5 | 0.5-3.5 |
| Pixel size (Å) | 0.73 | 0.73 | 0.73 |
| Symmetry imposed | C1 | C1 | C1 |
| Initial particle images (no.) | 3 376 077 | 3 376 077 | 3 376 077 |
| Final particle images (no.) | 98 911 | 98 911 | 98 911 |
| Map resolution (Å) | | 2.07 | 3.64 |
| FSC threshold | | 0.143 | 0.143 |
| Map resolution range (Å) | | ~1.8-2.2 | ~2.9-3.7 |
| **Refinement** | | | |
| Initial model (PDB code) | 9QTS | | |
| Model resolution (Å) | | | |
| FSC threshold | | | |
| Model resolution range (Å) | | | |
| Map sharpening *B* factor (Å$^2$) | | 36 | 60 |
| Model composition | | | |
| Non-hydrogen atoms | 27476 | | |
| Protein residues | 3394 | | |
| Ligands | 25 | | |
| *B* factors (Å$^2$) | | | |
| Protein | 119.93 | | |
| Ligand | 94.11 | | |
| R.m.s. deviations | | | |
| Bond lengths (Å) | 0.005 | | |
| Bond angles (°) | 0.735 | | |
| Validation | | | |
| MolProbity score | 1.74 | | |
| Clashscore | 13.89 | | |
| Poor rotamers (%) | 0.96 | | |
| Ramachandran plot | | | |
| Favored (%) | 97.55 | | |
| Allowed (%) | 2.42 | | |
| Disallowed (%) | 0.03 | | |
